# Bayestrat: Population Stratification Correction Using Bayesian Shrinkage Prior for Genetic Association Studies

**DOI:** 10.1101/2021.03.23.436705

**Authors:** Zilu Liu, Asuman Turkmen, Shili Lin

## Abstract

In genetic association studies with common diseases, population stratification is a major source of confounding. Principle component regression (PCR) and linear mixed model (LMM) are two commonly used approaches to account for population stratification. Previous studies have shown that LMM can be interpreted as including all principle components (PCs) as random-effect covariates. However, including all PCs in LMM may inflate type I error in some scenarios due to redundancy, while including only a few pre-selected PCs in PCR may fail to fully capture the genetic diversity. Here, we propose a statistical method under the Bayesian framework, Bayestrat, that utilizes appropriate shrinkage priors to shrink the effects of non- or minimally confounded PCs and improve the identification of highly confounded ones. Simulation results show that Bayestrat consistently achieves lower type I error rates yet higher power, especially when the number of PCs included in the model is large. We also apply our method to two real datasets, the Dallas Heart Studies (DHS) and the Multi-Ethnic Study of Atherosclerosis (MESA), and demonstrate the superiority of Bayestrat over commonly used methods.

## Introduction

There has been a long-standing interest to investigate associations between common diseases and single-nucleotide polymorphisms (SNPs) in genome-wide association studies (GWAS). Many of these investigations are population-based cohorts or case-control studies, wherein the main source of confounding arises from population stratification, corresponding to the systematic differences in allele frequencies between subpopulations due to ancestry^[1]^. It is well known that failure to appropriately account for population stratification may result in identifications of biologically irrelevant variants, and consequently, inflated type I error rates in statistical tests^[1–5]^.

Numerous statistical methods have been proposed to reduce the impact of population stratification in GWAS, including genomic control^[6,7]^, structured association^[8]^, adjustment by principal components (PCs)^[9–12]^, and mixed models^[13–16]^. Genomic control adjusts the test statistics by an overall inflation factor. Structured association divides samples into strata, and results are then combined after association analysis is performed within each stratum. Previous studies have shown that genomic control and structured association may not be effective in certain situations^[13,17]^. Therefore, recent attention has been focused on PCR and LMM. In PCR, population stratification is adjusted by including a few top PCs computed from null markers as covariates in regression models. As top PCs potentially represent ancestry or spatial information, PCR has been commonly used and shown to be effective in controlling population stratification^[11,12]^. However, prespecifying the optimal number of PCs to be included in the model so that one can sufficiently account for population stratification is a challenge. It is typical to include the top 5, 10 or 20 PCs in a regression model^[18–21]^. Nevertheless, it should be noted that a few top PCs may not represent population substructures, as they could simply be reflections of long-range linkage disequilibrium (LD) or assay artefacts^[10]^. Further, when more subtle and regional substructures exist, a few top PCs may not be sufficient to eliminate spurious associations caused by population stratification, in which case, a larger number of PCs might be needed^[22,23]^. Mathieson et al. ^[23]^ simulated a scenario where 100 PCs largely mitigated the inflated p-values, which was not able to be achieved by 20 PCs or fewer. Several PC selection methods have also been proposed, providing some guidance on which PCs should be included as covariates. These include the Tracy-Widom test that assesses the significance of eigenvalues^[11]^, selection of PCs by reduction of genomic control inflation factor^[24]^, the permutation-based test PC-Finder^[25]^ and selection of PCs that are significantly associated with the outcome^[26,27]^. Abegaz et al. ^[18]^ also suggested a forward or stepwise PC selection algorithm, which allows for automated variable selection as another way to determine which PCs should be included. One common drawback of all these methods is the requirement of a benchmark to decide which PCs should be kept in the model.

An alternative approach to PCR is mixed models, where a random effect, with an estimated genetic relationship matrix (GRM) as the covariance structure up to a scale, is included in the regression model^[16]^. Typically, a GRM is estimated empirically from a large number of null markers or known genealogies. The linear mixed model (LMM) has been commonly used to prevent false positive associations due to sample structures, including population structure and family relatedness, and has also been shown to be effective in increasing power in studies without sample structures^[28]^. Previous research has revealed the connections between LMM and PCR by showing that both of them essentially share the same underlying regression model when the Balding-Nichols GRM is considered^[18,29,30]^. The discrepancy between them is that LMM includes all PCs as random effects, whereas PCR only incorporates only a few selected PCs as fixed effects in the model. Nevertheless, LMM may cause an overfitting problem due to inclusion of redundant information that can dilute the influence of relevant PCs and degrade the quality of the correction^[29]^. On the other hand, an underfitting problem may arise from the traditional PCR approach given that population structures may not be fully captured by a few top PCs. As such, these overfitting and underfitting issues need to be investigated and addressed, despite the efficacies of PCR and LMM over genomic controls and structured association in controlling population stratification.

In this paper, we propose an approach under the Bayesian framework with the goal of identifying SNPs that are significantly associated with a continuous outcome while accounting for population stratification. Motivated by the connection between LMM and PCR, and the practical difficulty of choosing the optimal PCs, we propose a PC self-selection approach, Bayestrat, which imposes appropriate shrinkage priors on the PC effects and serves as an automatic PC selection procedure. In essence, our method can be viewed as a comprise between LMM and PCR from a Bayesian perspective. We investigate the performance of our method through a comprehensive simulation study. We also apply Bayestrat to two real data analyses to demonstrate its superiority to commonly used methods.

## Results

### Synopsis of Bayestrat

In order to remedy the potential underfitting with only a few top PCs as in PCR while avoiding overfitting induced by including all PCs as in LMM, we propose Bayestrat, an alternative approach by including a substantial number of PCs but subjecting each to a judicious selection procedure. Specifically, our procedure is aimed to shrink the estimated effect sizes of non-confounded PCs towards zero while boosting the signals of highly confounded ones. ***U***nder the Bayesian framework, we assign a shrinkage prior to each PC effect (Methods section). Given that our method incorporates a large number, although not necessarily all, of the PCs, it is a comprise between LMM and the traditional PCR models, where the former includes all PCs without shrinkage while the latter only includes a few PCs that may not be sufficient.

To illustrate the similarity yet also contrasting the differences between the proposed Bayestrat and the existing PCR and LMM methodologies, we summarize the workflows of these three approaches in Fig. 1, which also serves as a visual illustration of our perspective that Bayestrat is a compromise between PCR and LMM. When the number of PCs added to the model is large (e.g., >20PCs), Bayestrat facilitates the detection of highly confounded PCs, which may not be necessarily the top ones associated with the largest eigenvalues since PCs are obtained in an unsupervised manner. Meanwhile, Bayestrat also shrinks the effects of non- or minimally confounded PCs towards zero. Therefore, Bayestrat is an association testing procedure that enables automatic selection of highly confounded PCs, and in the mean time, provides high power for detecting SNPs that are associated with the disease.

**Fig. 1.**
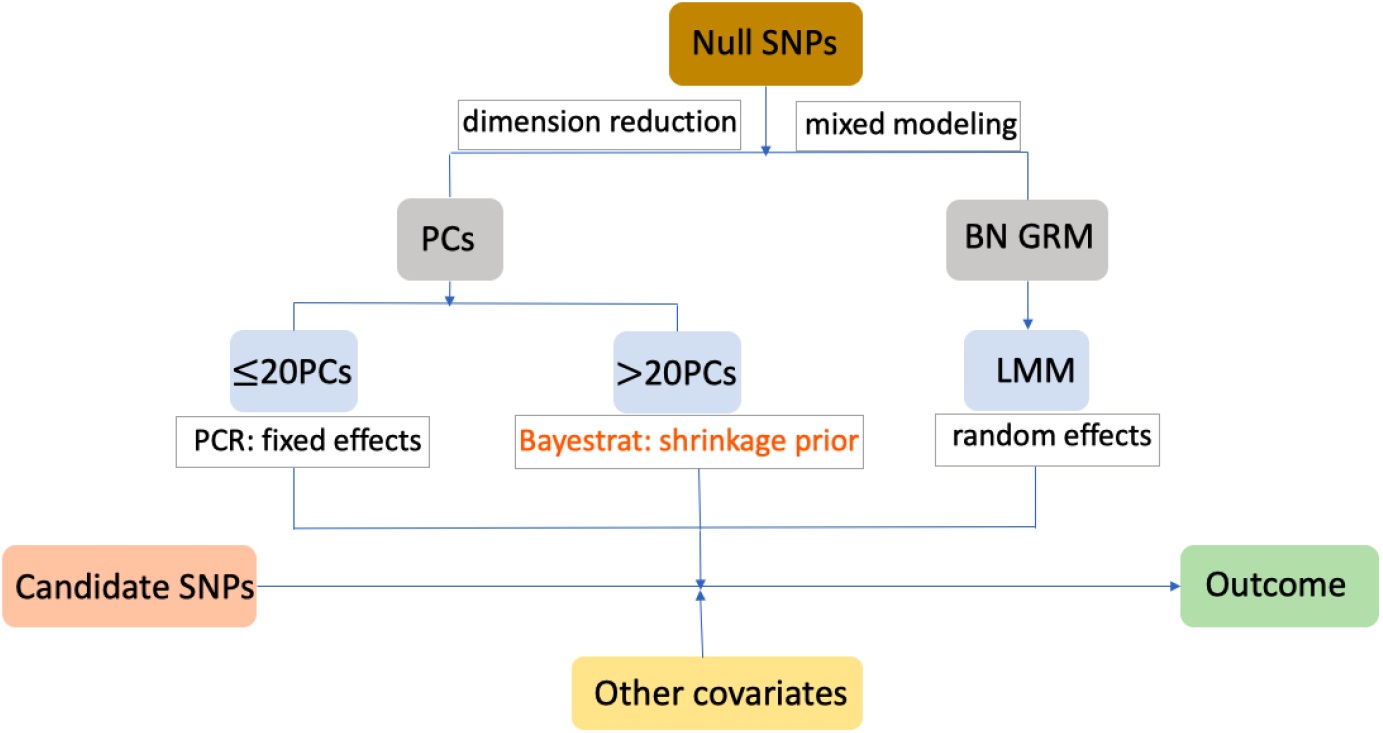
Similarities and differences among existing methods and the proposed Bayestrat method for addressing population stratification. PCs and the Balding-Nichols (BN) GRM are calculated from null SNPs. Two existing methods either include top (typically no more than 20) PCs as fixed-effect covariates (PCR), or include all PCs as random effects (LMM) for testing associations between candidate SNPs and a phenotype, after adjusting for other non-genetic covariates. We further note that LMM can be viewed as adding all PCs with normal priors from a Bayesian perspective. Our proposed Bayestrat includes a moderate to large (more than 20) number of PCs with shrinkage priors, which can be viewed as a compromise between PCR and LMM.

### Bayestrat achieves lower type I errors

We conduct a simulation study to investigate the performance of correcting for spurious SNP associations and controlling for type I errors under population stratification. We generate genotype data stratified by four populations using the Balding-Nichols model^[31]^. We consider four populations of equal size, with a total sample size of *n* = 200, 1000, or 2000 individuals. A total of 10100 ancestry allele frequencies are first generated, followed by generating the population-specific allele frequencies. The first 100 SNPs are reserved for association testing and the rest of the 10000 SNPs are treated as null markers for PCs and GRM computations. We consider three population structures (Struct 1, Struct 2, and Struct 3) to generate a continuous phenotype *Y*, which is assumed to be not associated with any SNPs for type I error investigation (Methods section). A total of nine methods are compared: without any adjustment, adjustment by the top 2 PCs as fixed effects, adjustment by the top 25, 200 or 400 PCs either with normal priors method or with Bayestrat (Laplace shrinkage priors), and adjustment by LMM.

For *n* = 1000, we observe that population stratification is a severe problem in all three population structures when no adjustment is made, exhibited by the highly inflated type I errors (Fig. 2a for Struct 1; Supplementary Figures S1a and S2a for Struct 2 and Struct 3, respectively). We can also see that only using the top two PCs are not sufficient to fully correct for spurious associations either, which is expected given that PC3 is an important confounder: it is correlated with the phenotype and also explains part of the ancestry background in all three structure models (Supplementary Table S1). Therefore, it is not always sufficient to include a few top PCs, especially when the optimal number of PCs cannot be predetermined. As more PCs are added (from 25 to 200 to 400), we can see that there is a slight trend of increased type I errors for the normal priors method, whereas Bayestrat (with Laplace shrinkage priors) is able to maintain acceptable error rates. This conclusion remains valid for the other two sizes (Supplementary Figure S3). Finally, LMM is seen to control spurious associations well for the majority of the SNPs but fails to correct for a number of false positives: highly inflated type I errors are seen in Struct 1 and Struct 2 (Fig 2a and Supplementary Figure S1a). Overall, Bayestrat is the only method that consistently controls type I errors among all three population structures considered.

**Fig. 2.**
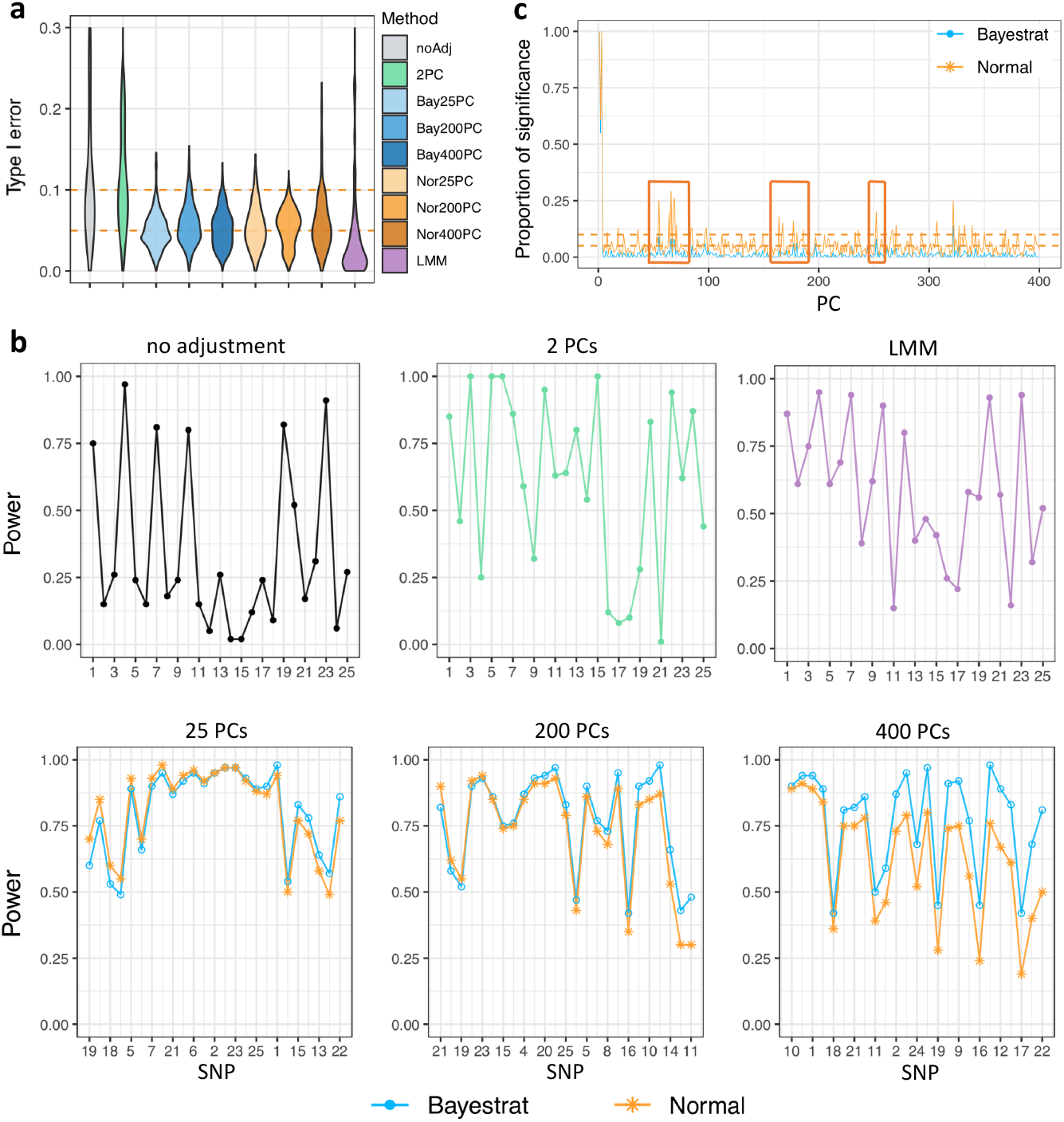
Type I error, power, and PC selection results for Struct 1 based on 100 replications. The total sample size is 1000 (250 for each population). the number of null SNPs is 10000 for obtaining PCs or GRM, and 100 SNPs are used for testing for associations. **a**. Type I errors for 100 SNPs set not to be associated with the trait value. The two horizontal dashed lines represent cutoffs of 0.05 and 0.1, respectively. Type I error is truncated at 0.3 for a better view. **b**. Power results for testing SNPs 1-25, set to be associated, from the power simulation study. For the last row, the x-axes are ordered by the power difference between Bayestrat and the normal priors method. **c**. Proportion of times, out of the 100 replications, the credible interval of a PC does not include 0 when testing for a spurious SNP with 400 PCs. The orange boxes capture sets of PCs where Bayestrat (with Laplace priors) shrinks down the detection of low or non-confounded PCs compared to the normal priors method.

### Bayestrat achieves higher power

To evaluate the power for detecting true SNP associations, we use the same data generation procedure, except that the first 25 out of the 100 reserved SNPs for association testing contribute to the phenotype. Effect sizes of the first 25 SNPs are set to be inversely proportional to their MAFs averaged over the four populations (Methods section). For testing the unassociated SNPs 26-100, we achieve the same conclusions as in the type I error study in the previous section (Supplementary Figure S4). For testing the associated SNPs 1-25, the power can be as low as zero when no adjustment for population structures is made (Fig. 2b). When adjusted by only two PCs, several SNPs are identified with high power, but some remain rarely detected. Bayestrat (with Laplace shrinkage priors) and the normal priors method both have high power when 25 or more PCs are included, although the advantage of Bayestrat is also clearly seen. The difference in the power grows larger as the number of PCs increases. This conclusion also holds for Struct 2 and Struct 3 (Supplementary Figure S1b and S2b). Since the sufficient number of PCs needed is unknown a priori; therefore, a safeguard against insufficient (underfitting) correction is to have a large enough number of PCs and let Bayestrat filter out those that are irrelevant. As expected, increasing the sample size leads to higher power in general for both types of priors (Supplementary Figure S5). Finally, LMM achieves competitive power in Struct 3 (Supplementary Figure S2b), but remains deficient for the other two population structures (Fig. S2b and Supplementary Figure S1b). Overall, Bayestrat is able to maintain high power among all three population structures considered when various numbers of PCs are included.

### Bayestrat facilitates accurate identification of highly confounded PCs

For the result of PC identifications in Struct 1, a commonality between Bayestrat and the normal priors method is that both achieve high proportion of detection for the top three PCs. The main difference is that some later PCs (Fig. 2c – enclosed in red boxes) are identified with substantial proportions by the normal priors method but are detected with relatively low frequencies using Bayestrat. As indicated by the negligible values of correlation with the phenotype, these later PCs contribute little to the phenotypic variance (Supplementary Table S1) and are unlikely to be important confounding factors in the association between the phenotype and the SNPs. As such, their detections are viewed as spurious, which may contribute to the worsening performance of the normal priors method for larger number of PCs. We observe similar PC selection results in Structure 2 (Supplementary Figure S1). In Struct 3, we observe that Bayestrat detects PC4 and those after less frequently than the normal prior model (Supplementary Figure S2). Although a total of 10 PCs are set to have non-zero effect sizes on the outcome, PC4 and those after have less contribution to the phenotypic variance (Supplementary Table S1), thus they are not the highly confounded factors in the association study. Overall, Bayestrat is able to capture highly confounded PCs and shrink the effects of less confounded or not confounded ones, resulting in more efficient PC identifications. Taken together with the type I error and power results, the ability of Bayestrat in shrinking irrelevant information improves the identification of real signals while keeping those that are unassociated at base.

### Analysis of the Dallas Heart Study Data

The Dallas Heart Study (DHS) sequencing data contains variants genotyped on three genes, ANGPTL3, ANGPTL4 and ANGPTL5 for four self-identified populations: non-Hispanic Black, non-Hispanic White, Hispanic, and others^[32,33]^. As amply discussed in the literature, use of discrete ancestry identifiers may not correctly account for population stratification as opposed to the use of continuous ancestry measures such as PCs^[34]^. Therefore, in this analysis, our goal is to identify SNPs that are associated with serum triglyceride (TG) levels by accounting for ancestry effect using PCs; the results from Bayestrat will also be compared with no adjustment or normal priors methods. For data preprocessing, we first apply a log-transformation on TG to address skewness. Individuals who were treated with statin or had missing statin status are removed from the data. The SNPs that have no sequence variation are also removed, resulting in 3236 individuals genotyped on 269 variants. Twenty four null SNPs with MAF> 0.01 are selected by the non-significance criteria based on PLINK^[35]^ association results (Methods section); they are then used for obtaining the PCs. Variable standardization is also performed, and age and sex are included as two biological covariates. Except the selected nulls, the rest of the SNPs are tested for association with log(TG) by three methods: the model without adjustment, Bayestrat with 24 PCs, and the model adjusted by 24 PCs with normal priors. However, we did not run LMM due to the nature of its computational intensiveness. Because of the candidate gene approach in the sequencing study, only a limited number of SNPs are identified as the nulls, and some may in fact be in high LD with the other SNPs. Despite this imperfect selection of nulls, the E40K variant in gene ANGPTL4 is detected by Bayestrat and the normal priors method as having a protective effect on the TG level (Supplementary Table S2). Consistent with our result, E40K was previously reported to be significantly associated with lower TG levels^[36,37]^. Variant Y386X in gene ANGPTL5 is selected by the model without adjustment and Bayestrat, but not the normal priors method. This SNP is a nonsense mutation and has been previously reported to be associated with TG in haplotype association studies^[38,39]^. On the other hand, all three methods detect M259T in gene ANGPTL3 with a protective effect. Romeo et al. ^[33]^ also reported the significance of this variant and substantiated the results using additional study data. Previous studies indicated that the M259T variant failed to suppress lipoprotein lipase activity in vitro, as compared with the wild type ANGPTL3 ^[33,40]^. Overall, Bayestrat is more powerful than both the normal priors or the no adjustment mothods. In our analyses, PC6 is detected as the only significant confounder among the 24 PCs with a positive effect size by both Bayestrat and the normal priors method (Supplementary Table S3).

### The Multi-Ethnic Study of Atherosclerosis

The Multi-Ethnic Study of Atherosclerosis (MESA) is an ongoing medical research study that is aimed to investigate the characteristics and predictive risk factors for subclinical cardiovascular diseases^[41]^. The sample consists of four self-reported ethnicities: Caucasian (42%), African-American (24%), Asian American (12%) and Hispanic (22%). After the first examination was conducted in 2002, three additional examinations were followed to assess clinical mortality. Various biological covariates were measured and SNPs were genotyped across the whole genome. We focus our analysis on the fourth main examination, and the HDL cholesterol (HDL-C) as the out-come variable. After deleting individuals with missing HDL-C or had no genotype data recorded, a total of 4832 subjects are left. As in the DHS analysis, log-transformation is applied to HDL-C to address the skewness in the data, and each covariate is standardized prior to the analysis, including age and sex.

### Genome wide analysis using PLINK for the MESA data

We first perform two whole genome association tests for the SNPs on the autosomes using PLINK, with the first one adjusting for age and sex (plink1), and the second one additionally adjusting for self-reported ethnicities (plink2) as a conventional way to correct for population stratification as a comparison (although not recommended^[34]^). A total of 508581 SNPs passing the criteria of p-value> 0.05 and MAF> 0.05 for both plink1 and plink2 are selected as null SNPs for calculating PCs. The first three PCs, together explaining 10.27% of the total SNPs variability, fail to provide a satisfactory separation of the four ethnic groups (Fig. 3a-b). Although the Asians are clustered together and separated from the rest, the three other populations are still tangled together. Compared to the other populations, the Hispanic is more diverse and mixed. Fig. 3a indicates that potential subgroups exist within the Hispanic population, which are obviously not differentiated by self-reported ethnicities nor the first three PCs. Given the complexity of genetic backgrounds, it is more reasonable to treat ancestry estimation as a continuous measure. As such, statistical adjustments based on Bayestrat or LMM would likely be more appropriate compared to self-reported ethnicities as a way to address population stratification. To be comprehensive, we also rerun PLINK by adding the first 20 PCs, in addition to age and sex, as fixed covariates (plink3). Without any adjustment for population stratification, plink1 identifies 5921 and 827 genome-wide significant SNPs for the nominal level of 10^−5^ and 5 ×10^−8^, respectively, out of the total 869224 tested (Fig. 3c inner ring; also see top segment of Table 1 for the numbers of SNPs). Adjustment by the top 20 PCs reduces the respective numbers to 1404 and 123 (Fig. 3c middle ring). On the other hand, adjustment by self-reported ethnicities appears to be overly conservative, leading to only 69 and 8 detected positives (Fig. 3c outer ring). The highly significant SNPs enclosed in the two orange boxes in Fig. 3c on chromosome 8 and 16 belong to gene LPL^[42–45]^ and LPCAT2 ^[46,47]^, respectively, which will be analyzed further in the next subsection.

**Fig. 3.**
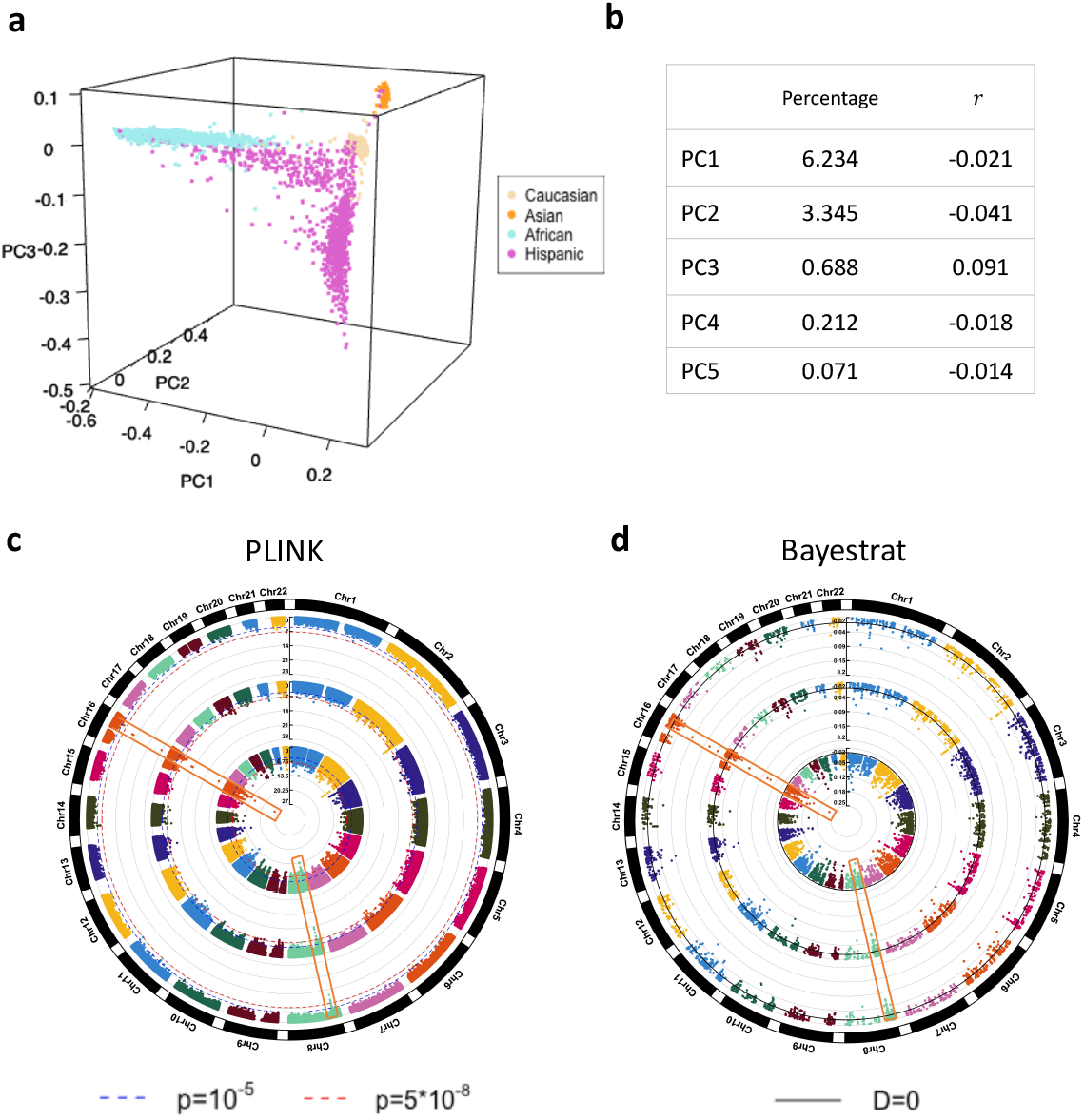
MESA data analysis results. **a**. 3-dimensional plot of the first three PCs, which are calculated from 508581 null SNPs. **b**. Percentage of total variability explained and *r*, Pearson correlation with log(HDL-C) for each of the first 5 PCs. **c**. Circular Manhattan plot for genome wide analysis using PLINK. The y-axis is −log_10_(p-value). Inner ring: plink1 (no adjustment for population stratification). Middle ring: plink3 (adjustment by top 20 PCs as covariates with fixed effects). Outer ring: plink2 (adjustment by self-reported ethnicities). **d**. Circular Manhattan plot for analyzing a filtered set of 5921 SNPs using Bayestrat, no adjustment, and normal priors methods, at a confidence level of 99.99%. The y-axis is a metric of significance, *D*, calculated from the credible interval (*L, U*) of a SNP effect. Specifically, *D* = *min*(*L, U*) if 0 ∉ (*L, U*); *D* = −*min*(*L, U*) if 0 ∈ (*L, U*). *D* > 0 indicates statistical significance (the larger the number, the more significant the result is). Inner ring: no PC included population stratification adjustment; Middle ring: Normal priors method adjusting for the top 200 PCs; Outer ring: Bayestrat adjusting for the top 200 PCs. The two orange boxes in both **c** and **d** capture the genomic region for LPL on chromosome 8 and that for LPCAT2 on chromosome 16.

**Table 1.**
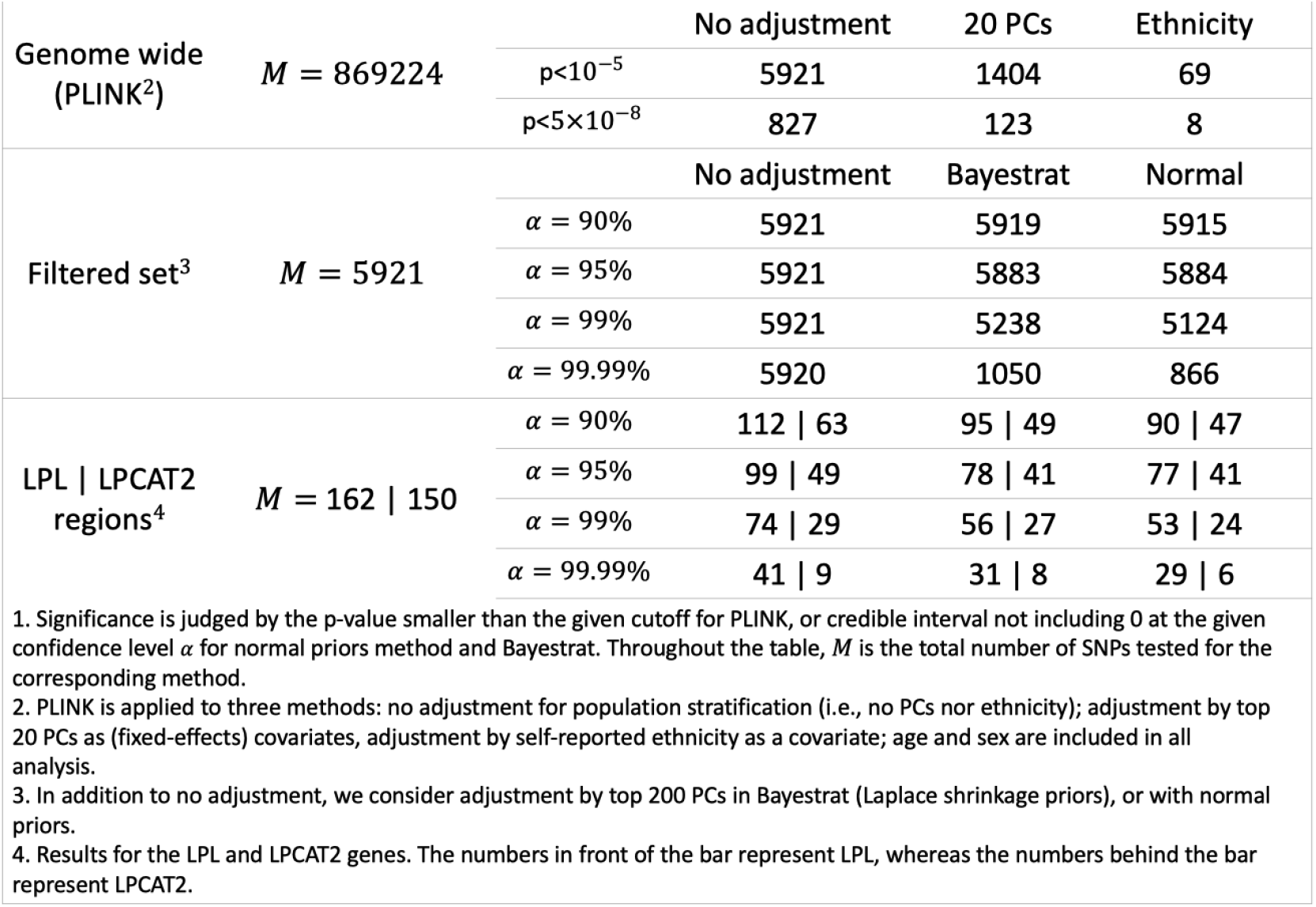
Number of significant SNPs detected by PLINK and Bayestrat^1^

### Bayestrat analysis on a filtered set of SNPs for the MESA data

The 5921 SNPs with p-value < 10^−5^ from plink1 are selected as candidate SNPs for Bayestrat analysis. We consider the method without adjustment (i.e. no PCs included), the normal priors method with the top 200 PCs, and Bayestrat (with Laplace shrinkage priors), also including 200 PCs. We again note that LMM is not performed due to computational cost. Confidence levels (α) of 90, 95, 99 and 99.99% are considered for credible interval calculations. As α increases from 90% to 99.99%, the number of detections changes slightly for the no adjustment method (middle segment of Table 1), whereas both the normal priors method and Bayestrat reduce the number of identifications substantially; further-more, Bayestrat identifies 21.25% more associated SNPs than the normal priors method for α = 99.99% (Fig. 3d, comparing the outer ring to the middle ring). Guided by the results from the simulation where Bayestrat was shown to be more powerful while controlling type I error, it is highly likely that the additional SNPs identified by Bayestrat are true positives. Consistent with the PLINK results, several SNPs in the LPL and LPCAT gene regions produce highly significant signals (Fig. 3d, orange boxes). Effect estimates for PCs are summarized in Supplementary Table S4.

### Analyses of LPL and LPCAT2 gene regions

To follow up on previously identified high-signal loci, we conduct further analysis for two gene regions, bp19775655 to bp20128692 for LPL on chromosome 8 (162 SNPs), and bp55644700 to bp55197000 for LPCAT2 on chromosome 16 (150 SNPs). The LPL gene provides instructions for making enzyme lipoprotein lipase, which plays an important role in breaking down fat in the form of triglycerides^[48]^. The LPCAT2 is a protein coding gene, which encodes a member of the lysophospholipid acyltransferase family. The encoded protein may function in membrane biogenesis and production of platelet-activating factor in inflammatory cells^[46]^. To investigate potential effects of these two genes on HDL-C, we analyze the data using the same three methods (no adjustment, normal priors method and Bayestrat). When testing LPL SNPs including 200 PCs for α = 99.99%, the normal priors method and Bayestrat reduce the number of significant SNPs that are detected with the no adjustment method (bottom segment of Table 1). It is particularly interesting to see that a group of four SNPs (Fig. 4, enclosed in the small gray circle on the bottom-right corner) detected by the no adjustment method are no longer significant in the two methods with PC adjustment. Another noteworthy pattern is that the cluster of 25 SNPs (Fig. 4, enclosed in the large gray circle) detected by at least one of the three methods are in high LD (Fig. 4, heatmap). In addition, the no adjustment method generally produces the highest absolute scaled effect sizes, defined as the mean divided by the standard deviation among the posterior samples of the corresponding effect. For SNPs that are significant in both the normal priors method and Bayestrat, the latter detects them with larger absolute scaled effect sizes. Although the differences are usually small, they are consistent for all SNPs, indicating greater power of Bayestrat. Moreover, three SNPs identified by Bayestrat are not significant in the normal priors method; they are rs1313432491 (bp19871869), rs17410962 (bp19892360) and rs1400899529 (bp19985425). Previous study has reported a significant association between rs17410962 and HDL^[43]^. On the other hand, rs1400899529 is located between two nearby expression quantitative trait loci (eQTL), rs113988682(bp19985257) and rs17091891(bp19985660), reported in GTEx^[49]^ of whole blood, spleen and nervet. Further note that the region flanked by these two eQTLs (indicated by the right thin vertical orange bar in Fig. 4) is contained within the high LD region, which provides evidence that rs1400899529 is a true positive. Similarly, rs1313432491 is within a high LD region flanked by two eQTLs, rs11986182 (bp19869936) and rs74549553 (bp19874863) (indicated by the left thick vertical orange bar in Fig. 4). All these provide further indication, with support from the literature, that Bayestrat is more powerful than its normal priors counterpart. Finally, two SNPs that are detected by all three methods with large scaled effect sizes, rs326 and rs327, were previously reported to be associated with the plasma lipid level^[45,50]^. Similarly, for LPCAT2 SNPs, adding 200 PCs helps to avoid three spurious findings from the no adjustment model at α = 99.99% (Supplementary Figure S6); furthermore, Bayestrat detects two more SNPs than the normal priors method. Finally, rs9989419 (bp55542640), commonly detected by normal priors method and Bayestrat, was previously reported to be associated with HDL-C^[45]^. Detailed information for the detected SNPs is summarized in Supplementary Tables S5 and S6.

**Fig. 4.**
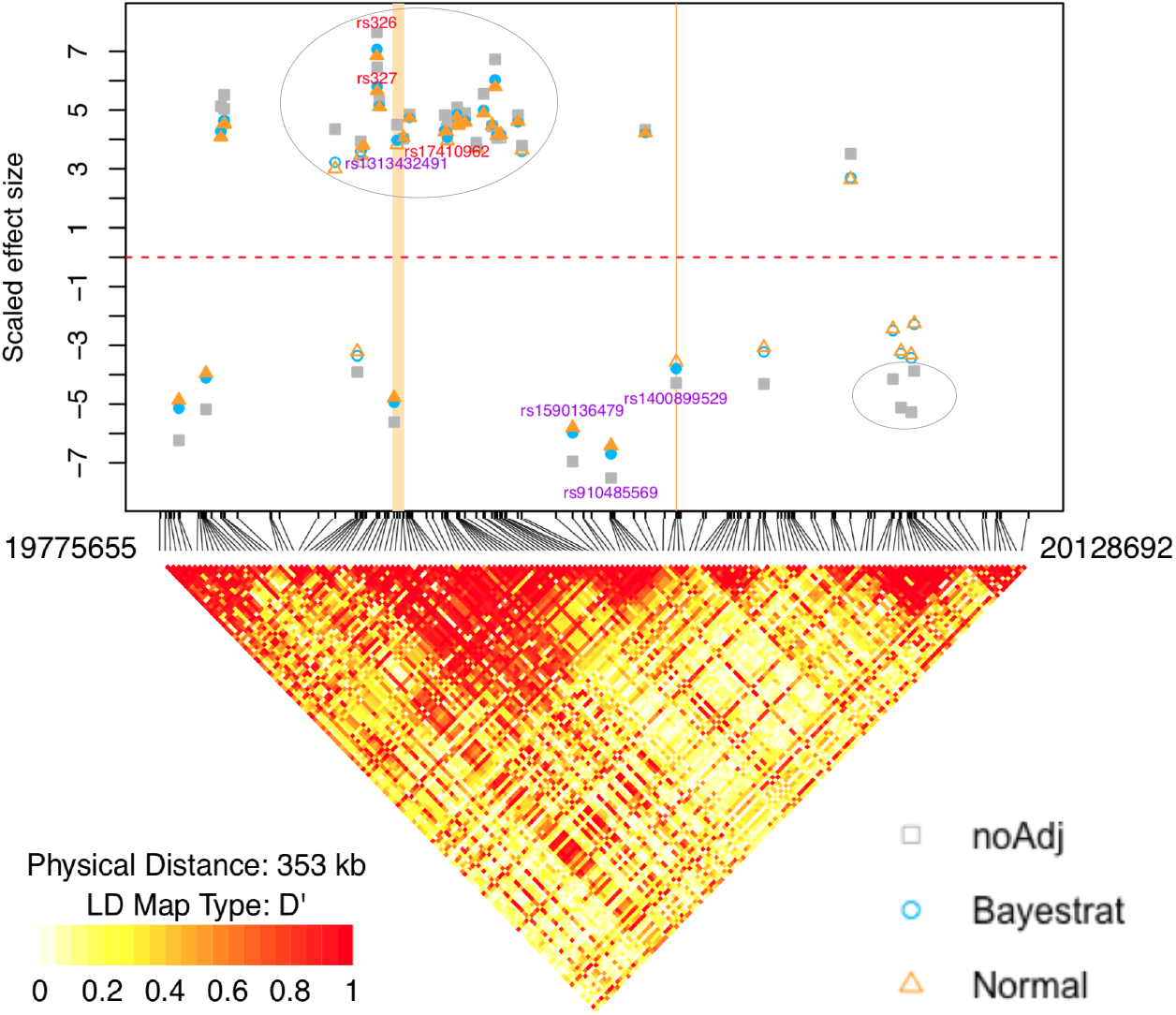
Scaled effect sizes and LD map of SNPs in a region containing gene LPL. Scaled effect size is the posterior mean divided by the posterior standard deviation. The scatter plot on the top of this figure presents the scaled effect sizes of SNPs that are detected by at least one of the three methods: no adjustment method, normal priors method, and Bayestrat. Colors are filled in for SNPs that are significant at α = 99.99%. The small grey circle on the right bottom encloses four SNPs believed to be spurious. The large grey circle on the upper left encloses a cluster of significant SNPs that are in high LD. The left thick vertical orange bar represents the region spanned by two reported eQTLs, rs11986182 and rs74549553. Similarly, the right thin vertical orange bar represents the region spanned by eQTL rs113988682 and rs17091891. The SNPs with rs number in red have been previously reported in the literature.

## Discussions

In this paper, we propose a statistical method, Bayestrat, to control for population stratification in population-based genetic association studies for continuous traits. Our method utilizes appropriate priors to implement a shrinkage scheme on PC effects, which overcomes the deficiencies of traditional PCR and LMM approaches. More specifically, the PCR approach with only a few leading PCs may be insufficient to fully capture the genetic background information in some cases, leading to underfitting. On the other hand, the popular mixed model approach has the deficiency of overfitting in our examples, due to the inclusion of a large amount of irrelevant noisy information. The proposed approach of including a large number of PCs assigned with shrinkage priors is essentially a compromise between these two approaches, which avoids the problem of underfitting by including a large number of PCs, yet remedy the overfitting problem by “eliminating” the effects of non- or minimally confounded PCs through automatic feature selection with shrinkage priors.

In simulation studies, we construct settings of population stratification based on population-specific means and individual PC scores. The results suggest that Bayestrat outperforms the non-shrinkage normal priors method, both for controlling type I error and enhancing power for detecting real associations, especially for the scenarios with a large number of PCs, owing to its shrinking effects on non-confounded PCs. Compared to adding a few top PCs and LMM, Bayestrat is able to significantly reduce spurious signals while maintaining high power consistently for the three population structure settings studied.

In the analysis of the DHS dataset, the no adjustment method and Bayestrat identify one SNP that was reported to be significant in haplotype association studies, which is not detected by the normal priors method. Another two previously reported associations are also identified in our analysis. However, we note that the analysis and interpretation are limited due to the small number of available null SNPs. For the MESA data analysis, adjustment by a sufficiently large number of PCs reduces the number of likely spurious associations to a large extent. On the other hand, the results also indicate that Bayestrat identifies more likely true associations and also detects SNPs with higher absolute effect sizes than the use of non-shrinkage normal priors. Some of the identified SNPs are reported in the literature or in high LD with those reported.

Bayestrat can be easily generalized to discrete outcomes by using a different modeling scheme. The logistic and probit models are common choices to model case-control status; the proportional odds model is suitable for ordinal traits. In addition, variable selection for PC effects can also be performed under the non-Bayesian framework, where automated PC selections are facilitated by imposing penalties on PC coefficients, with appropriate parameters to control the degrees of penalties^[51–54]^. A two-step analysis that selects PCs by a shrinkage scheme first and then conducts further analysis with only the selected PCs is also plausible.

## Methods

### The PCR approach

Let ***X*** be an *n* by *K* matrix of genotype scores for *n* individuals on *K* null markers, and let ***X***^∗^ be standardized version of ***X***, where 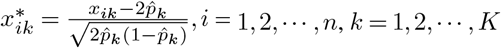 and 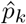 is the estimated frequency of the minor allele from the data. Consider the singular value decomposition (SVD) for ***X***^∗^ such that ***X***^∗^ = ***U****SV* ^*T*^, where ***U*** is an *n* by *n* matrix of left singular vectors (the columns), *S* is an *n* by *K* rectangular diagonal matrix of singular values, and *V* is a *K* by *K* matrix of right singular vectors (the columns). ***U***tilizing the SVD, PCs are obtained by *R* = ***X***^∗^*V* = ***U****S*, where *R* = (*R*_1_,, *R*_*n*_)*T* represents an *n* by *L* matrix of PC scores, *L* = *min* {*n, K*}

Let *y* = (*y*_1_, *y*_2_,, *y*_*n*_)^*T*^ be a quantitative trait vector for *n* subjects, and *g* = (*g*_1_, *g*_2_,, *g*_*n*_)^*T*^ be a vector of genotype scores, where *g*_*i*_ takes values in {0,1,2} representing the number of minor alleles individual *i* has on a specific SNP, *i* = 1, 2,, *n*. Let *Z* = (*Z*_1_, *Z*_2_,, *Z*_*n*_)^*T*^ be a matrix of non-genetic covariates, such as age and sex. The PCR method includes PCs as covariates with fixed effects when modeling the relationship between *y, g* and *Z* as follows:

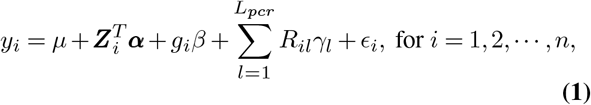

where *µ* is the intercept, **α**, *β* and ***γ*** = (*γ*_1_, *γ*_2_, …, *γL*_*pcr*_) are the effects of covariates, the genetic variant, and the PCs, respectively, and *L*_*pcr*_ is the number of preselected top PCs, typically no more than the top 20 PCs^[18]^. In addition, 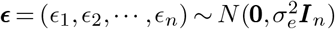 represents the collection of random error terms, where ***I***_*n*_ is the identity matrix of size *n*.

### The LMM approach

In LMM, a Genetic Relationship Matrix (GRM), denoted as an *n* by *n* matrix Φ, is introduced to measure genetic similarities between pairs of individuals, which is typically estimated from the null marker matrix ***X***. Two commonly used measures are the identity-by-state (IBS) matrix and the Balding-Nichols (BN) marix^[55,56]^. The IBS matrix measures the proportion of alleles identical by state between each pair of participants, i.e., 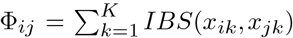 for individuals *i* and *j*, where *IBS*(*x*_*ik*_, *x*_*jk*_) ∈{0, 1, 2}represents the number of alleles that are of the same type between these two individuals. The BN matrix measures genetic similarities by 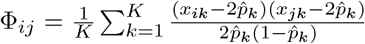. We consider the BN matrix in this manuscript to achieve equivalency between the PCR (when *L*_*P CR*_ = *L*) and the LMM approaches. The LMM considers both fixed and random effects in the regression model^[16]^,

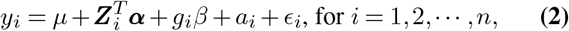

where the random effect 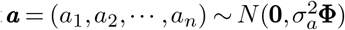 is used to correct for population stratification through incorporating Φ to serve as the covariance structure.

The equivalence between PCR and LMM holds due to the property of multivariate Gaussian distribution, and the following factorization of Φ:

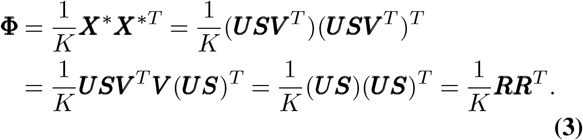

Now assume the PC effects 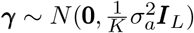. then it follows that 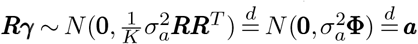 where 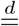 denotes equivalence in distribution. Therefore, the LMM essentially models all PCs as random-effect covariates, whereas the PCR approach typically includes a few preselected PCs into the model as fixed-effect covariates, thus can be viewed as a dimension-reduced approximation to LMM

### The proposed Bayestrat approach

Since PCR typically in-cludes only a few top PCs corresponding to the largest eigen-values, which may lead to underfitting (i.e., not being able to capture all population substructure), whereas LMM may have the opposite problem of overfitting due to including irrelevant PCs, we propose Bayestrat under the Bayesian setting as a compromise. Our model is,

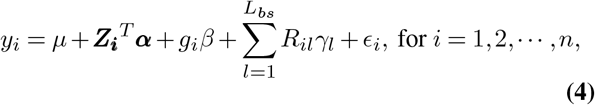

where *L*_*bs*_ is the number of top PCs to be fitted by Bayestrat, which is typically chosen to be much larger than *Lpcr* in PCR but much smaller than all PCs as in LMM.

In (4), the *ϵ*_*i*_’s are assigned independent and identically distributed (i.i.d.) normal priors, i.e.,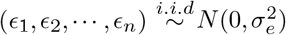, and 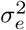 is assigned an Inverse-Gamma prior *IG*(*a*_*e*_, *b*_*e*_) for conjugacy. In order to shrink the effects of the PCs, we impose a Laplace prior with prior variance 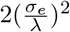 for each *γ*_*l*_ ^[57]^:

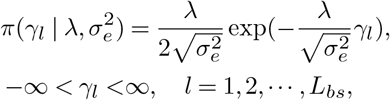

where λ is a hyperparameter contributing to the degree of penalty. We let λ^2^ follow *Gamma*(*a*_λ_, *b*_λ_) with density function 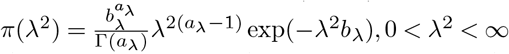. For the other parameters *µ*, **α** and *β* in (4), we impose independent ***U***niform priors on the real line, effectively assuming a fixed effect for each of them. For comparison purpose, we also consider normal priors as opposed to Laplace priors, leading to what we referred to as the normal priors method. Specifically, we set 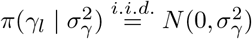, and 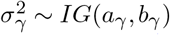. We follow the general guideline provided by Perez et al. ^[58]^ to specify parameters *a*_*e*_, *b*_*e*_, *a*_λ_, *b*_λ_, *a*_*γ*_ and *b*_*γ*_.

To achieve comparability among variables, standardization is performed on the outcome, PCs and other non-SNP covariates before analysis. Specifically, *y*_*i*_ is standardized by 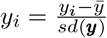, where 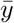 is the mean of the outcome and *sd*(*y*) is the standard deviation of the outcome variable. The PCs are standardized to be comparable to the SNP by 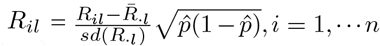 and *l*=1,…*L*_*bs*_ where 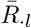 and *sd*(*R*_·*l*_) are the mean and standard deviation of the *l*^*th*^ PC, and 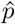 is the estimated minor allele frequency. The non-SNP covariates are standardized in the same way as the PCs.

### Computational consideration and vimplementation

Given the null marker matrix ***X***, we calculate the PCs and the GRM by R functions *prcomp* ^[59]^ or *flashpca* ^[60]^ for medium- and large-sized data (*n* × *K* ≤ 107). For even larger datasets such as MESA, we use the command *pca* in PLINK^[35]^ for the sake of computational speed. Bayestrat adapts much of the implemented Markov chain Monte Carlo (MCMC) algorithm in the R package BLR (Bayesian Linear Regression)^[58,61]^, which is designed for genomic selection and genetic values prediction using dense molecular markers. Significance is judged by the credible intervals of posterior samples not including zero under the given confidence level. Convergence diagnostics are conducted by the Raftery dependence factor and the Gelman-Rubin reduction factor^[62–64]^.

### Simulation studies

Let *pA*_*k*_ denote the ancestry allele frequency for SNP *k* and *pS*_*jk*_ the population-specific allele frequency for SNP *k* in population *j*. Based on the Balding-Nichols model^[31]^, we generate allele frequencies for both candidate and null SNPs in four populations by letting *p A*_*k*_ ∼ ***U*** (0.05, 0.95), *k* = 1, 2, …, 10100 and 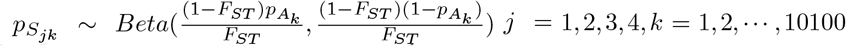. The Wright’s fixation index, *F*_*ST*_^[65]^, controling the degree of population differentiation, is set to be *F*_*ST*_ = 0.01. If *p S* _*jk*_> 0.5, we reset *pS*_*jk*_ = 1 − *pS*_*jk*_ to represent minor allele frequency. A continuous phenotype *Y* is then generated under the following three settings of population structures:

Struct 1: A mean-shift model with random error, that is

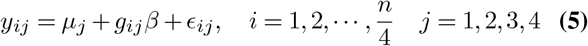

where *n* is the total number of subjects, and thus the number of subjects in each population is *n/*4. The mean shift parameter for each population is set to be *µ*_*j*_ = 0, 3, 6, 9 for *j* = 1, 2, 3, 4, respectively. In addition, 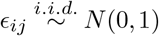.

Struct 2: This is also a mean-shift model, but the *µ*_j_ for each population is set to be 0, 0, 0, 5 for *j* = 1, 2, 3, 4, respectively This model only sets the fourth population apart from the others, as opposed to all four separate populations in Struct 1. Struct 3: This is a direct PC model with random errors by setting

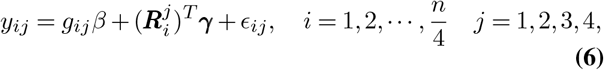

Where 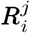 is a vector of PC scores for individual *I* in population *j*. We select 10 PCs among the first 50 to have non-zero effect sizes, and denoting them as *PC*_*l*_, *l* ∈ *A* = {1, 2, 3, 6, 14, 16, 22, 31, 37, 46}. Let *s*_*l*_ represent its corresponding singular value. To set the effect sizes for the selected PCs, we first calculate *b* × *s*_*l*_, *l* ∈ *A*, where *b* is chosen such that *b* × *s*_1_ = 0.05. These 10 numbers are then reordered and assigned as effect sizes *γ*_*l*_, *l* ∈ *A*, for PCs in the index set *A*: *γ*_*A*_ = {0.0489, 0.0271, 0.0500, 0.0268, 0.0268, 0.0497, 0.0266,0.0264,0.0263,0.0261}. The effect sizes for all other PCs (*l* ∉ *A*) are set to be zeros. Although this setting may be viewed as matching the PCR or the LMM methods, not all PCs play a role nor all PCs playing a role belong only to the top few.

For the type I error study, we set *β* = 0 for each test SNP. For the power study, we set 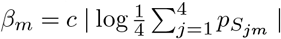 for *m* = 1, 2,, 25 and *β*_*m*_ = 0 for *m* = 26, 27,, 100, where *c* is chosen to be 0.15 to balance the effect size of genetic and confounding effects. By setting the non-zero *β*’s this way, we can see that the effect size gets larger as the average minor allele frequencies across the four populations gets smaller. In terms of computing the GRM, we set 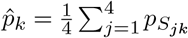. The prior distributions are set to be λ^2^ ∼ *G*(1.01, 0.01), 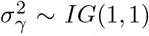, and 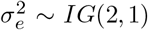 for our analysis. We also conduct a sensitivity analysis to investigate the impact of various priors, including 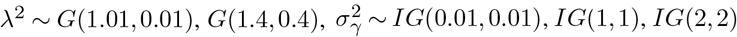, and 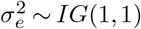. Results show that our simulation is not very sensitive to the selection of the priors considered here (Supplementary Figure S7-S9).

### Real data analysis

For the DHS data, the null SNPs are selected as those with MAF> 0.01 and p-value> 0.05*/*269 using PLINK. Due to the limited null information, we set a smaller MAF and p-value threshold (compared to the MESA analysis), and we also manually add three other SNPs, ANG3-005506-IVS4-137, ANGPTL-@8429-IVS632, and ANGPTL5-12766-IVS5-57, to the null set. These three SNPs are added because they are (almost) not significant based on the results from haplotype association studies (results are not presented in this manuscript for brevity). The rest of the 245 SNPs are tested for association with the standardized log(*T G*). We use the same hyperparameter settings as in the simulation study. For the MESA data, the prior distributions are set to be, 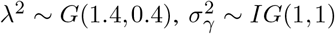, and 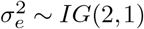. We use a different prior for λ as we would like to demonstrate the use of a variety of priors since our sensitivity study in the simulation shows that the results are comparable using different priors.

## Data availability

The simulated data used in this manuscript are provided on Github (https://github.com/Zilu-Liu/Simulation/tree/main/Bayestrat). The DHS dataset is accessible at https://www.utsouthwestern.edu/research/translational-medicine/doing-research/dallas-heart/. The MESA dataset is available at dbGap (https://www.ncbi.nlm.nih.gov/projects/gap/cgi-bin/study.cgi?study_id=phs000209.v2.p1) under accession numbers pht001120.v10.p3 and phg000071.v2 for phenotype trait tables and genotype datasets, respectively. The R package Bayestrat is freely available on Github (https://github.com/Zilu-Liu/Bayestrat).

## Acknowledgements

The authors would like to acknowledge the National Heart, Lung, and Blood Institute (NHLBI) that supports MESA data providers. Support for MESA is provided by contracts N01-HC95159, N01-HC-95160, N01-HC-95161, N01-HC-95162, N01-HC-95163, N01-HC-95164, N01-HC-95165, N01-HC95166, N01-HC-95167, N01-HC-95168, N01-HC-95169 and CTSA ***U***L1-RR-024156; funding for genotyping was provided by contract N02-HL-64278. The authors are also grateful to the Hoffman Family Center in Genetics and Epidemiology and the National Center for Advancing Translational Sciences (NCATS) for supporting the DHS study.

## Author contributions

S.L. designed the study. A.T. coordinated the project. Z.L. conducted the research. S.L. supervised the project. All authors discussed the results and contributed to the manuscript writing.

## Additional information

Correspondence and requests for materials should be addressed to S.L.

**Supplementary Figure S1.**
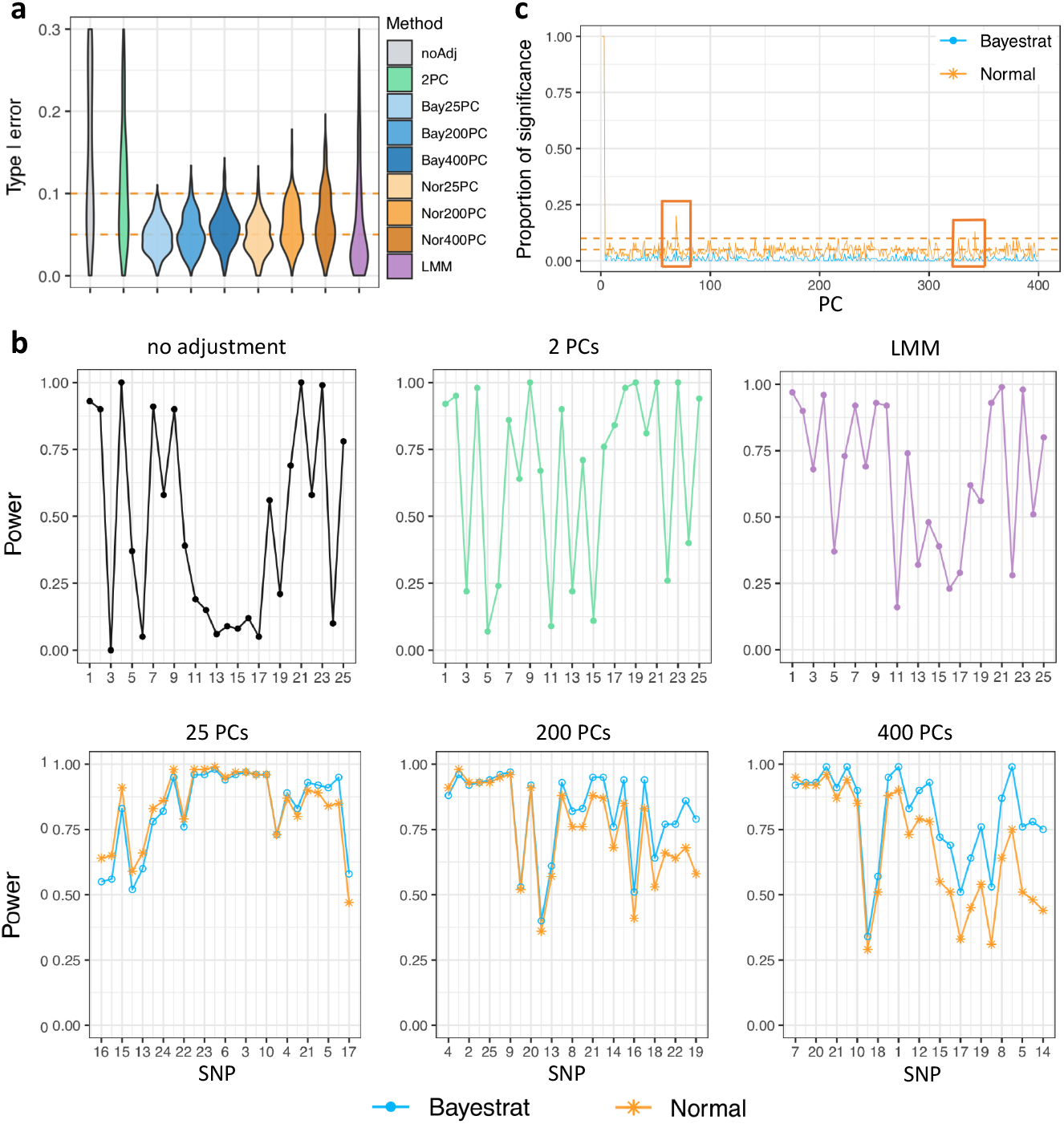
Type I error, power, and PC selection results for Struct 2 based on 100 replications. The total sample size is 1000 (250 for each population). the number of null SNPs is 10000 for obtaining PCs or GRM, and 100 SNPs are used for testing for associations. **a**. Type I errors for 100 SNPs set not to be associated with the trait value. The two horizontal dashed lines represent cutoffs of 0.05 and 0.1, respectively. Type I error is truncated at 0.3 for a better view. **b**. Power results for testing SNPs 1-25, set to be associated, from the power simulation study. For the last row, the x-axes are ordered by the power difference between Bayestrat and the normal priors method. **c**. Proportion of times, out of the 100 replications, the credible interval of a PC does not include 0 when testing for a spurious SNP with 400 PCs. The orange boxes capture sets of PCs where Bayestrat (with Laplace priors) shrinks down the detection of low or non-confounded PCs compared to the normal priors method.

**Supplementary Figure S2.**
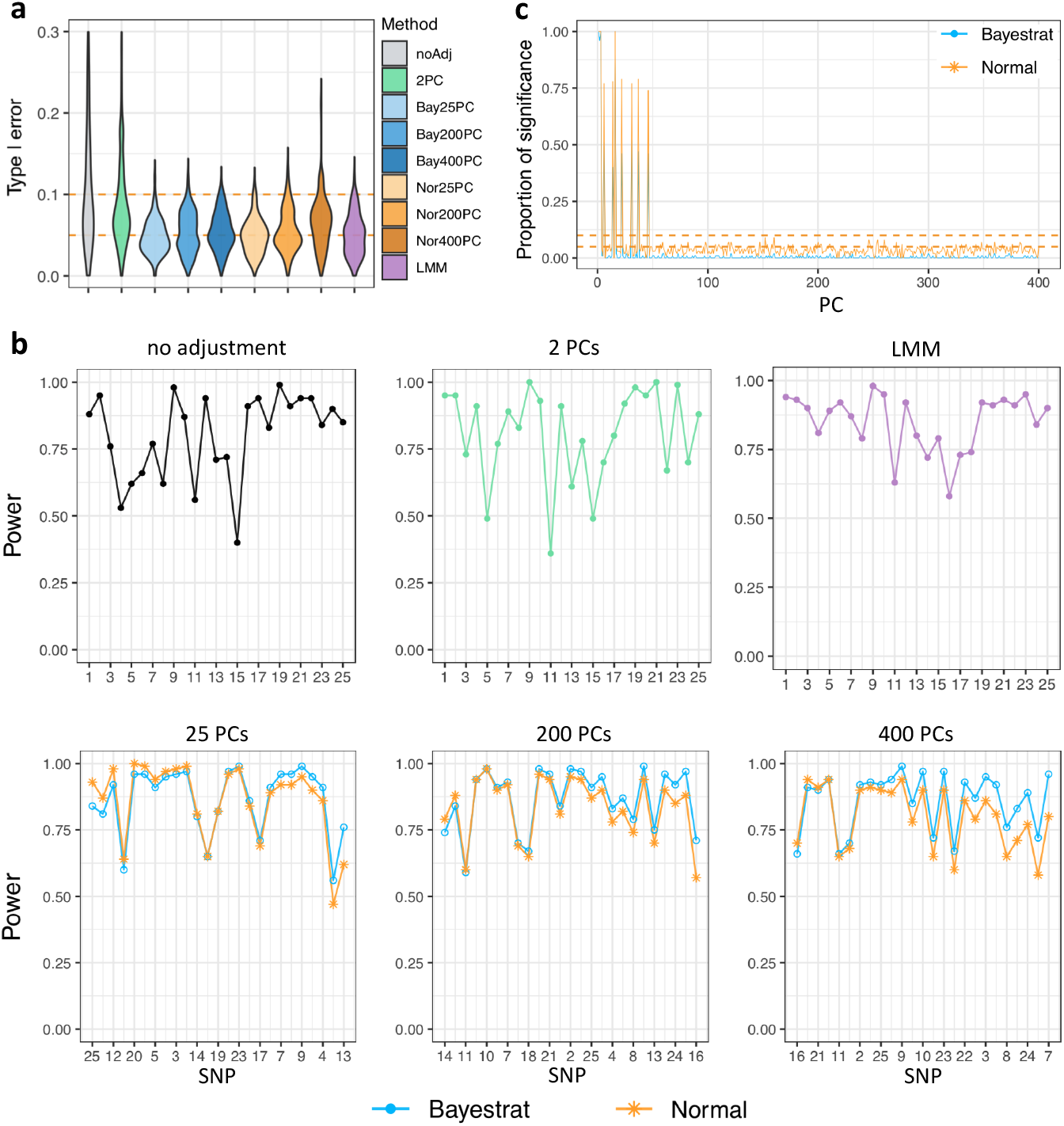
Type I error, power, and PC selection results for Struct 3 based on 100 replications. The total sample size is 1000 (250 for each population). the number of null SNPs is 10000 for obtaining PCs or GRM, and 100 SNPs are used for testing for associations. **a**. Type I errors for 100 SNPs set not to be associated with the trait value. The two horizontal dashed lines represent cutoffs of 0.05 and 0.1, respectively. Type I error is truncated at 0.3 for a better view. **b**. Power results for testing SNPs 1-25, set to be associated, from the power simulation study. For the last row, the x-axes are ordered by the power difference between Bayestrat and the normal priors method. **c**. Proportion of times, out of the 100 replications, the credible interval of a PC does not include 0 when testing for a spurious SNP with 400 PCs.

**Supplementary Figure S3.**
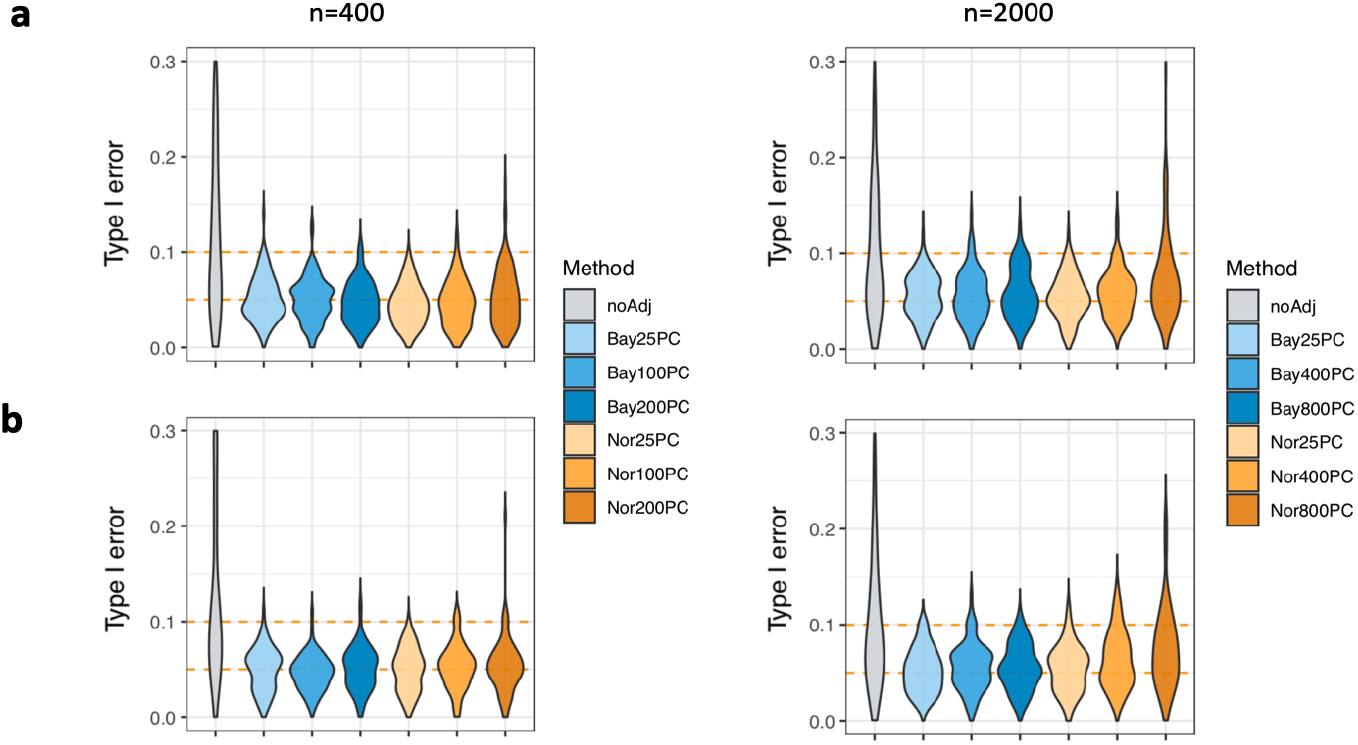
Type I error results with various sample sizes for Struct 1. *n* is the total number of subjects for four equal-sized populations. **a**. Results for all 100 SNPs from the null model. **b**. Results for SNPs 26-100 from the power simulation study.

**Supplementary Figure S4.**
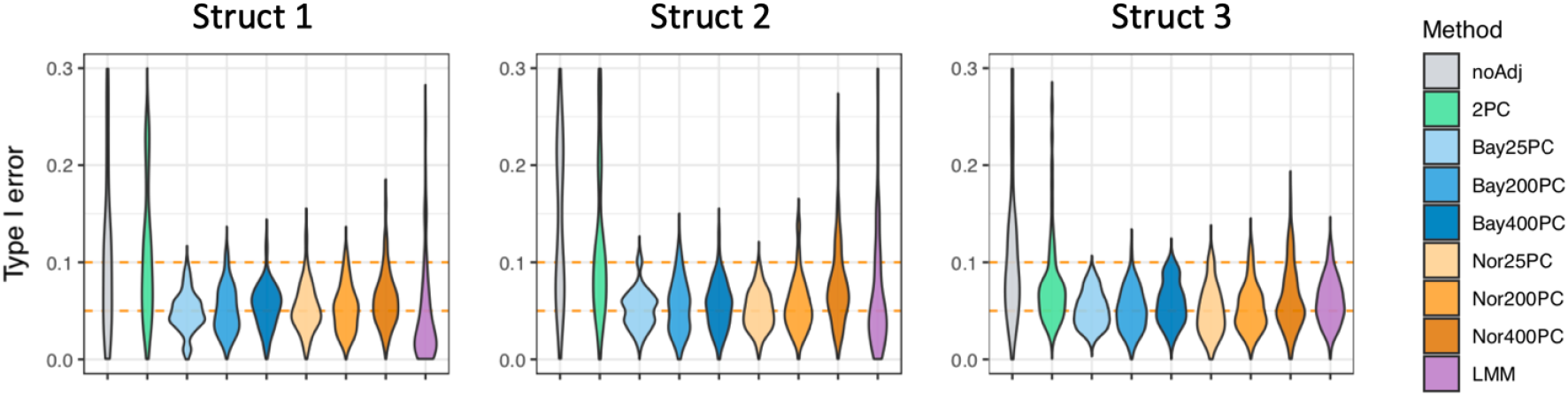
Type I error results for SNPs 26-100 from the power simulation study for three population structures with total sample size 1000.

**Supplementary Figure S5.**
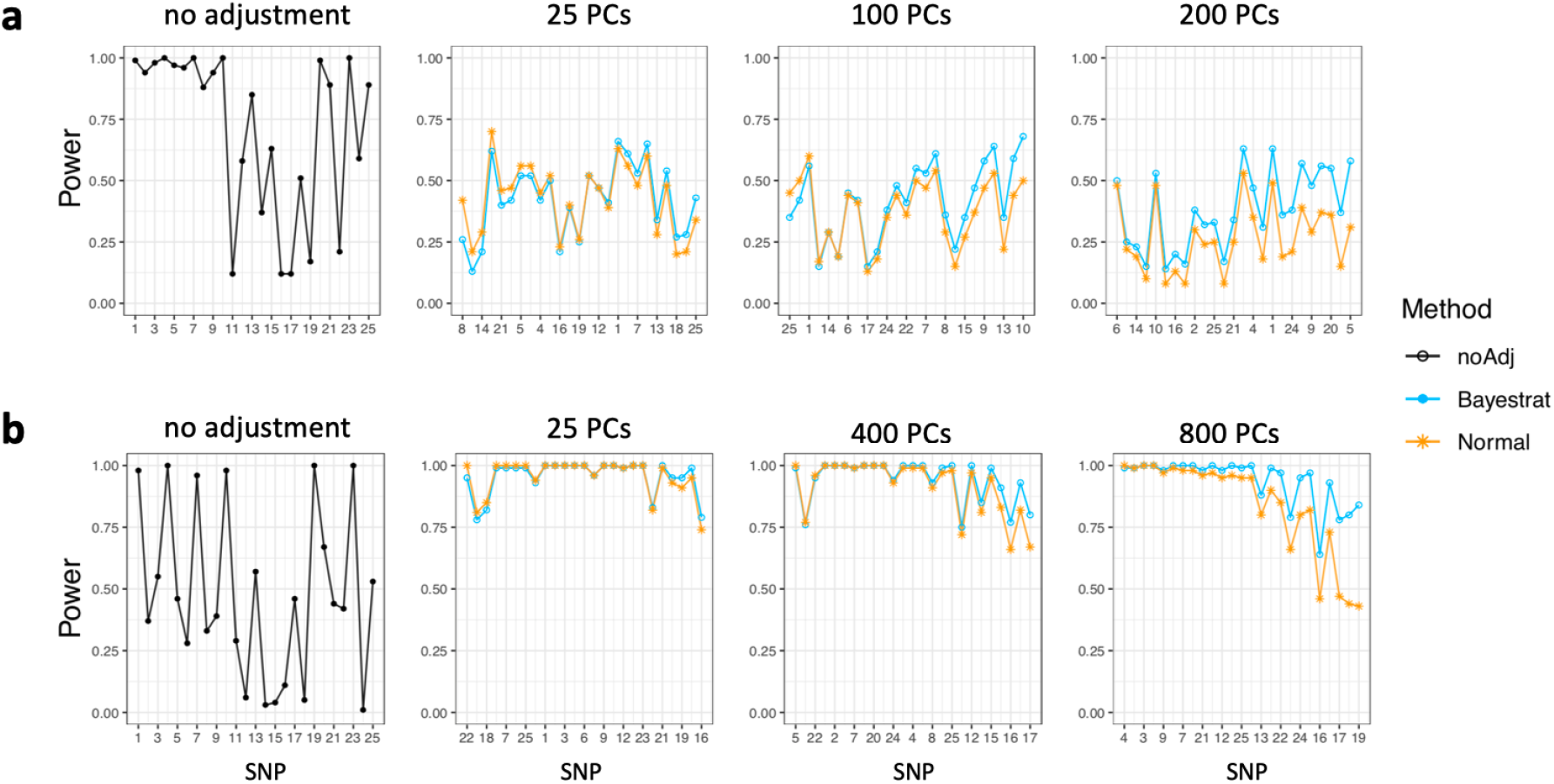
Power results with various sample sizes in simulation Struct 1. **a**. The total sample size *n* = 400. **b**. *n* = 2000. For columns other than no adjustment, x-axes are ordered by power difference between Bayestrat and the Bayesian model with normal priors.

**Supplementary Figure S6.**
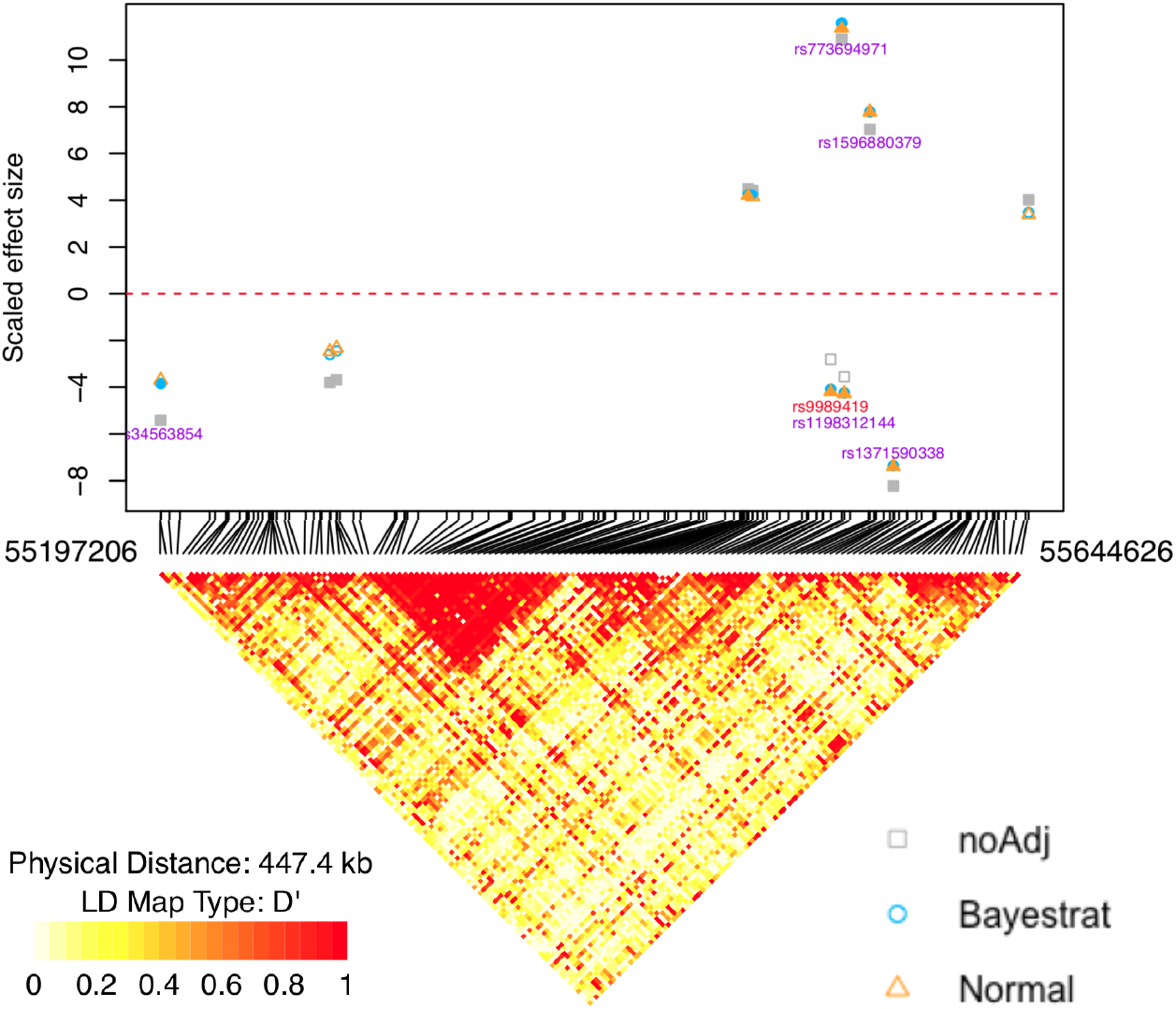
Scaled effect sizes and LD map of SNPs in a region containing gene LPCAT2. Scaled effect size is the posterior mean divided by the posterior standard deviation. The scatter plot on the top of this figure presents the scaled effect sizes of SNPs that are detected by at least one of the three methods: no adjustment method, normal priors method, and Bayestrat. Colors are filled in for SNPs that are significant at α = 99.99%. The SNPs with rs number in red have been previously reported in the literature.

**Supplementary Figure S7.**
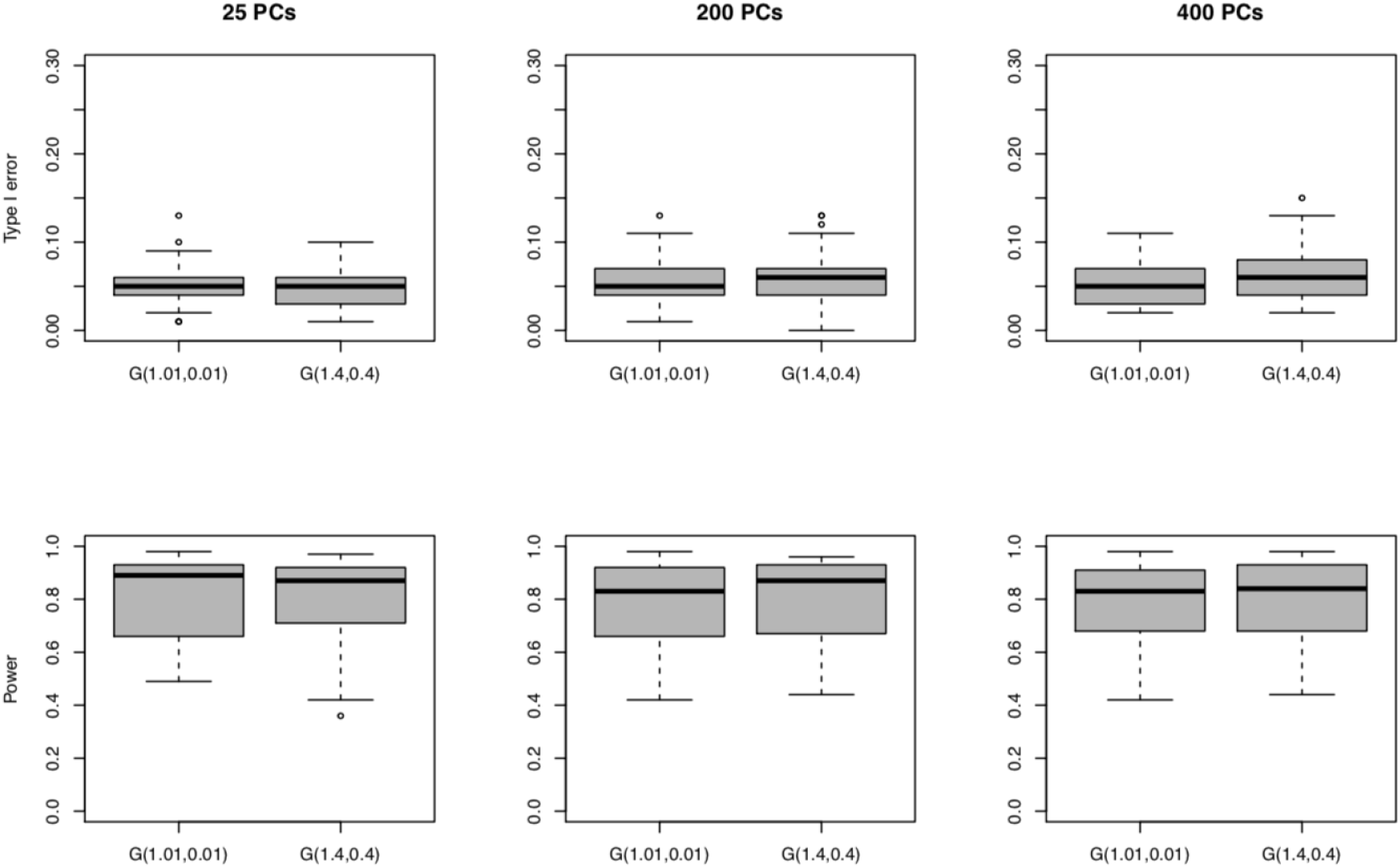
Sensitivity study for λ^2^ in simulation Struct 1. Consider λ^2^ ∼ *G*(1.01, 0.01) or *G*(1.4, 0.4). Type I error results are from null models. Power results are for SNPs 1-25 from power simulation studies.

**Supplementary Figure S8.**
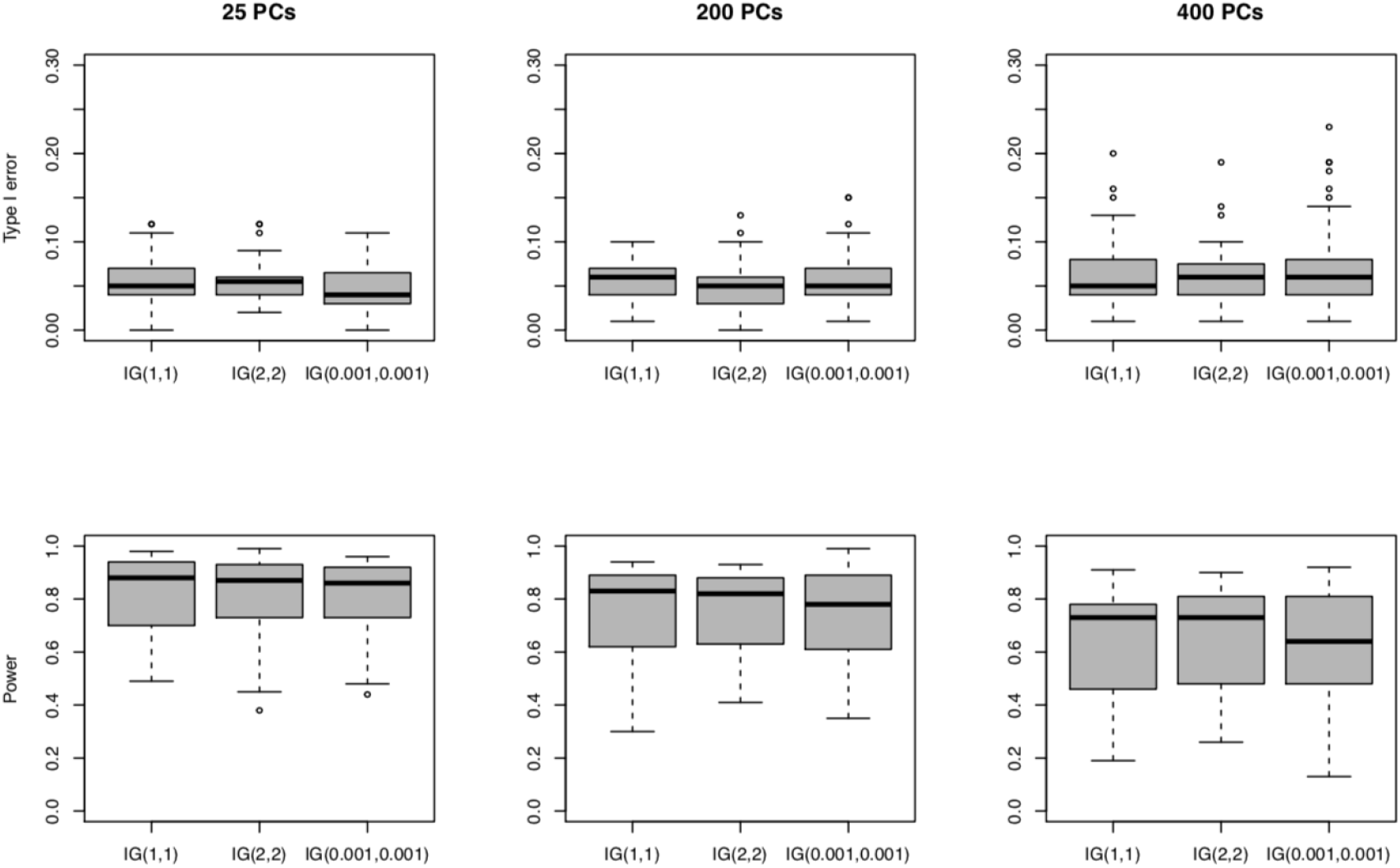
Sensitivity study for 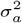 in simulation Struct 1. Consider 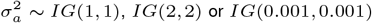. Type I error results are from null models. Power results are for SNPs 1-25 from power simulation studies.

**Supplementary Figure S9.**
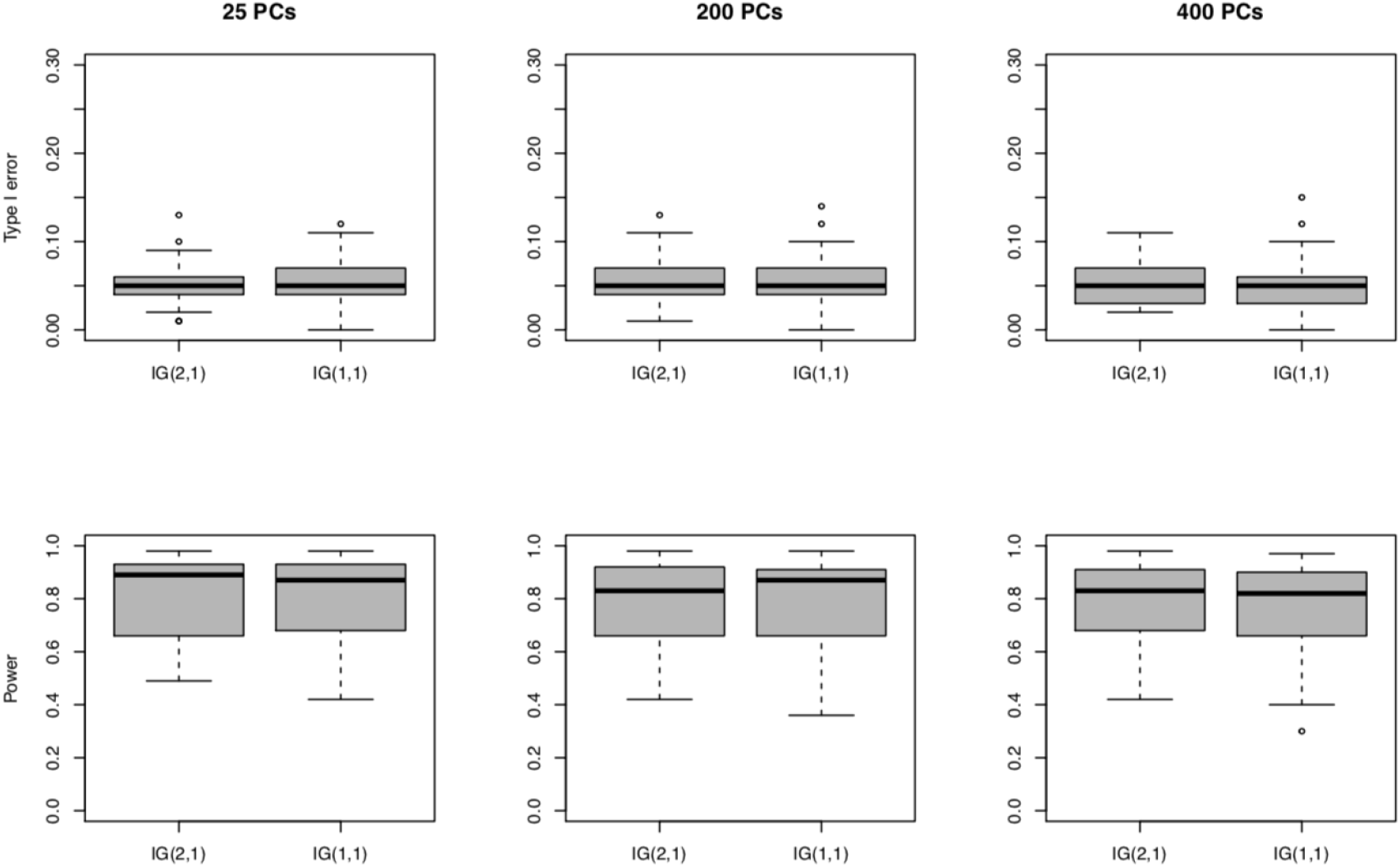
Sensitivity study for 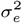 for Laplace PCs in simulation Struct 1. Consider 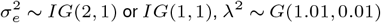. Type I error results are from null models. Power results are for SNPs 1-25 from power simulation studies.

**Supplementary Table S1.** Proportion of total variability explained by PCs and Pearson correlation (*r*) with the outcome in three simulation structures for sample size 1000. 1. Percentage of total variability of the null dataset explained by each PC. Regardless of different simulation settings, these proportions are unchanged. For Struct 1 and 2, correlations up to the top 20 PCs are reported. For Struct 3, correlations for the top 20 PCs and those selected to have non-zero effect sizes in the simulation model are reported.

| Name | Percentage<sup>1</sup> | $r_1$ | $r_2$ | $r_3$ |
| --- | --- | --- | --- | --- |
| PC1 | 0.572 | -0.915 | -0.799 | 0.296 |
| PC2 | 0.565 | -0.025 | -0.194 | 0.170 |
| PC3 | 0.547 | -0.233 | 0.350 | 0.329 |
| PC4 | 0.170 | -0.011 | -0.013 | -0.019 |
| PC5 | 0.169 | -0.004 | 0.004 | -0.004 |
| PC6 | 0.168 | 0.003 | 0.009 | 0.090 |
| PC7 | 0.168 | 0.005 | 0.016 | 0.032 |
| PC8 | 0.167 | 0.014 | 0.014 | 0.011 |
| PC9 | 0.167 | -0.016 | -0.027 | -0.052 |
| PC10 | 0.166 | -0.006 | -0.006 | -0.012 |
| PC11 | 0.166 | -0.007 | -0.013 | -0.038 |
| PC12 | 0.166 | 0.000 | -0.009 | -0.015 |
| PC13 | 0.165 | 0.010 | 0.009 | 0.009 |
| PC14 | 0.165 | -0.013 | -0.018 | 0.062 |
| PC15 | 0.165 | -0.002 | 0.001 | 0.011 |
| PC16 | 0.164 | 0.001 | -0.001 | 0.174 |
| PC17 | 0.163 | -0.008 | -0.008 | -0.017 |
| PC18 | 0.163 | -0.015 | -0.011 | -0.013 |
| PC19 | 0.163 | 0.009 | 0.013 | 0.031 |
| PC20 | 0.163 | 0.033 | 0.041 | 0.080 |
| PC22 | 0.162 |  |  | 0.064 |
| PC31 | 0.159 |  |  | 0.055 |
| PC37 | 0.158 |  |  | 0.072 |
| PC46 | 0.156 |  |  | 0.089 |

**Supplementary Table S2.** Credible intervals for significant SNPs detected by at least one of the three methods for DHS analysis (* for not significant). Significance is judged by 95% CI not including 0.

| SNP | Mutation type | MAF | noAdj |  | Bayestrat |  | Normal |  |
| --- | --- | --- | --- | --- | --- | --- | --- | --- |
|  |  |  | LB | UB | LB | UB | LB | UB |
| 005308-M259T | MISSEN | 0.0234 | -0.586 | -0.273 | -0.569 | -0.251 | -0.563 | -0.248 |
| 005645-IVS4-127 | INTRONIC | 0.0002 | 0.537 | 4.362 | 0.531 | 4.378 | 0.548 | 4.336 |
| 8357-non-coding | NONCODING | 0.3749 | -0.166 | -0.072 | -0.167 | -0.071 | -0.167 | -0.073 |
| @1175-Intronic | INTRONIC | 0.0387 | -0.429 | -0.19 | -0.416 | -0.17 | -0.411 | -0.168 |
| @1313-E40K | MISSEN | 0.0074 | * | * | -0.595 | -0.036 | -0.597 | -0.036 |
| @1318-M41I | MISSEN | 0.0042 | 0.162 | 0.904 | 0.136 | 0.878 | 0.132 | 0.869 |
| @6020-IVS373 | INTRONIC | 0.0002 | 0.972 | 4.825 | 0.933 | 4.744 | 0.936 | 4.764 |
| @6052-IVS341 | INTRONIC | 0.0262 | -0.526 | -0.236 | -0.509 | -0.216 | -0.502 | -0.211 |
| @6175-P210P | SYN | 0.0443 | -0.389 | -0.163 | -0.375 | -0.143 | -0.367 | -0.136 |
| @6219-IVS412 | INTRONIC | 0.0002 | 0.263 | 4.103 | 0.168 | 3.978 | 0.17 | 3.991 |
| @7870-IVS461 | INTRONIC | 0.0002 | 0.022 | 3.858 | 0.139 | 3.986 | 0.179 | 4.009 |
| @8191-R278Q | MISSEN | 0.0283 | -0.555 | -0.276 | -0.54 | -0.256 | -0.532 | -0.253 |
| @8279-P307P | SYN | 0.0618 | -0.361 | -0.169 | -0.354 | -0.155 | -0.344 | -0.145 |
| @8280-V308M | MISSEN | 0.0005 | 0.553 | 2.767 | 0.577 | 2.769 | 0.582 | 2.791 |
| @10620-N360N | SYN | 0.0418 | -0.389 | -0.167 | -0.375 | -0.151 | -0.37 | -0.146 |
| @10707-P389P | SYN | 0.0632 | 0.148 | 0.337 | 0.139 | 0.331 | 0.138 | 0.331 |
| @10769-Intronic | INTRONIC | 0.0643 | -0.334 | -0.152 | -0.323 | -0.134 | -0.318 | -0.131 |
| 26238-Y386X | NONSENSE | 0.0039 | -0.875 | -0.115 | -0.97 | -0.021 | * | * |

**Supplementary Table S3.** Coefficient estimates and credible intervals for 24 PCs for the DHS data. Estimations are averaged over all tested SNPs and all replications, and transformed back to the original scale. α = 95%.

| Name | Bayestrat |  |  | Normal |  |  |
| --- | --- | --- | --- | --- | --- | --- |
|  | Mean | LB | UB | Mean | LB | UB |
| PC1 | -0.042 | -0.11 | 0.026 | -0.042 | -0.114 | 0.029 |
| PC2 | -0.074 | -0.16 | 0.012 | -0.072 | -0.153 | 0.009 |
| PC3 | -0.007 | -0.075 | 0.061 | -0.008 | -0.089 | 0.073 |
| PC4 | -0.057 | -0.142 | 0.028 | -0.058 | -0.147 | 0.031 |
| PC5 | 0.072 | -0.025 | 0.169 | 0.072 | -0.026 | 0.171 |
| PC6 | 0.198 | 0.009 | 0.386 | 0.187 | 0.053 | 0.321 |
| PC7 | 0.018 | -0.072 | 0.108 | 0.02 | -0.088 | 0.127 |
| PC8 | -0.022 | -0.114 | 0.07 | -0.024 | -0.132 | 0.085 |
| PC9 | -0.071 | -0.179 | 0.037 | -0.072 | -0.187 | 0.042 |
| PC10 | 0.008 | -0.084 | 0.101 | 0.009 | -0.102 | 0.12 |
| PC11 | -0.056 | -0.159 | 0.047 | -0.058 | -0.172 | 0.056 |
| PC12 | 0.052 | -0.051 | 0.154 | 0.054 | -0.061 | 0.169 |
| PC13 | 0.023 | -0.074 | 0.12 | 0.025 | -0.09 | 0.14 |
| PC14 | -0.016 | -0.114 | 0.082 | -0.017 | -0.134 | 0.1 |
| PC15 | -0.024 | -0.124 | 0.076 | -0.026 | -0.145 | 0.093 |
| PC16 | 0.024 | -0.079 | 0.126 | 0.025 | -0.097 | 0.147 |
| PC17 | -0.077 | -0.198 | 0.043 | -0.079 | -0.208 | 0.049 |
| PC18 | -0.112 | -0.255 | 0.032 | -0.112 | -0.255 | 0.031 |
| PC19 | 0.047 | -0.089 | 0.182 | 0.049 | -0.109 | 0.207 |
| PC20 | -0.104 | -0.3 | 0.092 | -0.108 | -0.325 | 0.109 |
| PC21 | 0.046 | -0.144 | 0.236 | 0.05 | -0.176 | 0.275 |
| PC22 | 0.108 | -0.223 | 0.439 | 0.116 | -0.272 | 0.503 |
| PC23 | -0.071 | -0.401 | 0.259 | -0.077 | -0.47 | 0.317 |
| PC24 | -0.115 | -0.547 | 0.317 | -0.123 | -0.635 | 0.39 |

**Supplementary Table S4.** Coefficient estimates and credible intervals for the first 10 PCs for testing 5921 MESA SNPs. Estimations are averaged over all tested SNPs and all replications, and transformed back to the original scale. α = 95%.

| Name | Bayestrat |  |  | Normal |  |  |
| --- | --- | --- | --- | --- | --- | --- |
|  | Mean | LB | UB | Mean | LB | UB |
| PC1 | -0.029 | -0.123 | 0.058 | -0.036 | -0.135 | 0.063 |
| PC2 | -0.075 | -0.212 | 0.05 | -0.088 | -0.228 | 0.052 |
| PC3 | 0.803 | 0.483 | 1.123 | 0.782 | 0.477 | 1.086 |
| PC4 | -0.372 | -0.905 | 0.106 | -0.438 | -0.964 | 0.088 |
| PC5 | -0.318 | -1.185 | 0.458 | -0.402 | -1.309 | 0.505 |
| PC6 | -0.858 | -1.926 | 0.118 | -0.988 | -2.03 | 0.055 |
| PC7 | 0.099 | -0.812 | 1.044 | 0.129 | -0.922 | 1.180 |
| PC8 | -0.208 | -1.189 | 0.703 | -0.27 | -1.334 | 0.794 |
| PC9 | 0.142 | -0.797 | 1.129 | 0.185 | -0.904 | 1.273 |
| PC10 | 0.14 | -0.814 | 1.143 | 0.182 | -0.925 | 1.289 |

**Supplementary Table S5.** Credible intervals for the significant SNPs on the gene LPL region detected by at least one of the listed methods with α = 99.99% (* for not significant). rs number is provided when available.

| Position | rs | MAF | noAdj |  | Bayesrat |  | Normal |  |
| --- | --- | --- | --- | --- | --- | --- | --- | --- |
|  |  |  | LB | UB | LB | UB | LB | UB |
| 19783125 | rs1563270328 | 0.427 | -0.184 | -0.042 | -0.178 | -0.026 | -0.177 | -0.024 |
| 19794163 |  | 0.126 | -0.243 | -0.044 | -0.238 | -0.007 | -0.236 | -0.002 |
| 19800257 |  | 0.192 | 0.031 | 0.212 | 0.014 | 0.203 | 0.008 | 0.202 |
| 19801580 | rs976533691 | 0.146 | 0.033 | 0.241 | 0.023 | 0.233 | 0.018 | 0.23 |
| 19801599 |  | 0.404 | 0.031 | 0.171 | 0.015 | 0.173 | 0.016 | 0.183 |
| 19846682 | rs1231420054 | 0.477 | 0.011 | 0.142 | * | * | * | * |
| 19855697 |  | 0.493 | -0.142 | 0 | * | * | * | * |
| 19857092 |  | 0.237 | 0.005 | 0.177 | * | * | * | * |
| 19857982 |  | 0.136 | * | * | * | * | 0.005 | 0.221 |
| 19863719 | rs326 | 0.364 | 0.075 | 0.218 | 0.065 | 0.224 | 0.067 | 0.228 |
| 19863816 | rs327 | 0.308 | 0.054 | 0.207 | 0.042 | 0.202 | 0.04 | 0.206 |
| 19864713 |  | 0.099 | 0.055 | 0.292 | 0.041 | 0.286 | 0.049 | 0.291 |
| 19870653 |  | 0.418 | -0.182 | -0.034 | -0.175 | -0.026 | -0.176 | -0.021 |
| 19871869 | rs1313432491 | 0.069 | 0.031 | 0.269 | 0.011 | 0.294 | * | * |
| 19874450 |  | 0.062 | 0.013 | 0.312 | 0.016 | 0.32 | 0.002 | 0.325 |
| 19876926 |  | 0.091 | 0.037 | 0.281 | 0.026 | 0.283 | 0.031 | 0.298 |
| 19891340 |  | 0.121 | 0.031 | 0.254 | 0.013 | 0.244 | 0.016 | 0.246 |
| 19891970 | rs1234291694 | 0.087 | 0.014 | 0.289 | 0.013 | 0.282 | 0.02 | 0.283 |
| 19892360 | rs17410962 | 0.149 | 0.028 | 0.219 | 0.007 | 0.212 | * | * |
| 19896325 |  | 0.257 | 0.026 | 0.194 | 0.022 | 0.189 | 0.022 | 0.195 |
| 19896590 |  | 0.256 | 0.023 | 0.191 | 0.016 | 0.188 | 0.021 | 0.191 |
| 19896642 | rs1179382877 | 0.255 | 0.021 | 0.187 | 0.019 | 0.184 | 0.017 | 0.195 |
| 19896798 | rs993238185 | 0.256 | 0.025 | 0.194 | 0.015 | 0.188 | 0.005 | 0.187 |
| 19896866 |  | 0.256 | 0.024 | 0.184 | 0.014 | 0.185 | 0.017 | 0.189 |
| 19899552 |  | 0.257 | 0.014 | 0.194 | 0.018 | 0.187 | 0.016 | 0.188 |
| 19904071 | rs556303990 | 0.263 | 0.004 | 0.158 | * | * | * | * |
| 19907068 | rs73599626 | 0.349 | 0.033 | 0.189 | 0.026 | 0.178 | 0.016 | 0.18 |
| 19910554 | rs1035503181 | 0.23 | 0.008 | 0.183 | 0.016 | 0.193 | 0.014 | 0.197 |
| 19911725 |  | 0.395 | 0.056 | 0.193 | 0.044 | 0.21 | 0.046 | 0.204 |
| 19913956 | rs566154583 | 0.228 | * | * | 0.007 | 0.18 | 0.008 | 0.194 |
| 19912570 |  | 0.229 | 0.005 | 0.176 | 0.01 | 0.183 | 0.009 | 0.187 |
| 19921026 |  | 0.089 | 0.038 | 0.282 | 0.026 | 0.282 | 0.025 | 0.3 |
| 19922636 |  | 0.354 | 0 | 0.14 | * | * | * | * |
| 19943326 | rs1590136479 | 0.315 | -0.207 | -0.057 | -0.212 | -0.046 | -0.215 | -0.04 |
| 19958878 | rs910485569 | 0.346 | -0.215 | -0.065 | -0.217 | -0.058 | -0.218 | -0.056 |
| 19972862 |  | 0.111 | 0.015 | 0.246 | 0.016 | 0.24 | 0.021 | 0.247 |
| 19985425 | rs1400899529 | 0.359 | -0.151 | -0.009 | -0.159 | 0 | * | * |
| 20021096 | rs149480896 | 0.446 | -0.151 | -0.009 | * | * | * | * |
| 20056363 |  | 0.298 | 0.001 | 0.145 | * | * | * | * |
| 20073544 |  | 0.28 | -0.163 | -0.011 | * | * | * | * |
| 20076867 |  | 0.247 | -0.186 | -0.028 | * | * | * | * |
| 20080993 |  | 0.239 | -0.192 | -0.033 | * | * | * | * |
| 20082218 |  | 0.298 | -0.155 | -0.003 | * | * | * | * |

**Supplementary Table S6.**
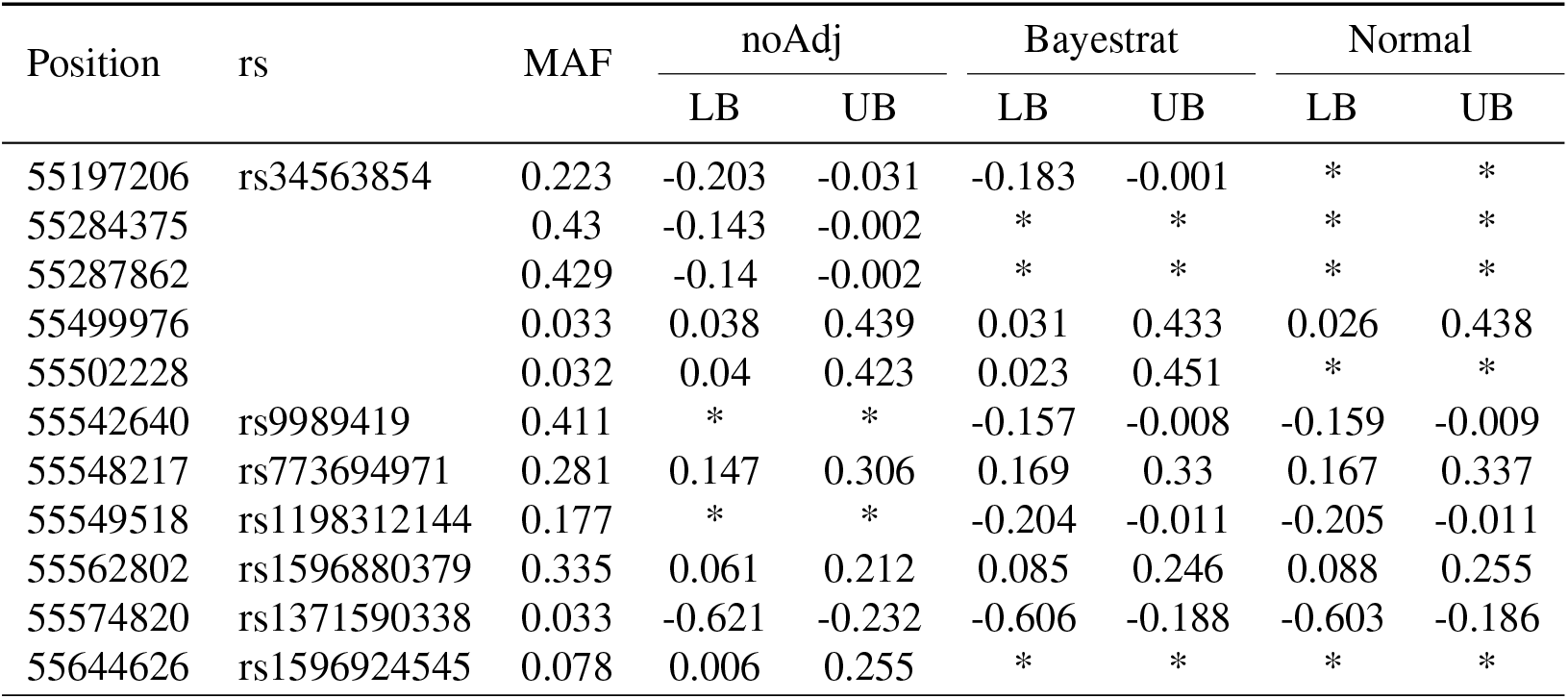
Credible intervals for the significant SNPs on the gene LPCAT2 region detected by at least one of the listed methods with α = 99.99% (* for not significant). rs number is provided when available.

